# Mismatch repair disturbs meiotic class I crossover control

**DOI:** 10.1101/480418

**Authors:** Tim J. Cooper, Margaret R. Crawford, Laura J. Hunt, Marie-Claude Marsolier-Kergoat, Bertrand Llorente, Matthew J. Neale

## Abstract

Sequence divergence, mediated by the anti-recombinogenic activity of mismatch repair (MMR), forms a barrier to meiotic recombination and in turn the formation of viable gametes. However, rather than MMR acting as a non-specific impediment to meiotic recombination, here we provide evidence that at regions of greater sequence divergence MMR preferentially suppresses interfering (class I) crossovers (COs). Specifically, as measured in two *Saccharomyces cerevisiae* hybrids containing thousands of DNA-sequence polymorphisms, removal of MMR components increases both the frequency of CO formation and the uniformity of the observed CO distribution. At fine scale, CO positions are skewed away from polymorphic regions in MMR-proficient cells, but, critically, not when members of the class I CO pathway, *MSH4* or *ZIP3*, are inactivated. These findings suggest that class I COs are more sensitive to heteroduplex DNA arising during recombination. Simulations and analysis of Zip3 foci on meiotic chromosomes support roles for Msh2 both early and late in the class I CO maturation process. Collectively, our observations highlight an unexpected interaction between DNA sequence divergence, MMR, and meiotic class I CO control, thereby intimately linking the regulation of CO numbers and their distribution to pathways contributing to reproductive isolation and eventual speciation.

## Introduction

Meiosis, a specialised two-step nuclear division, is responsible for the generation of genetically diverse, haploid gametes. An integral feature of the meiotic program is the initiation of homologous recombination via programmed Spo11-dependent DNA double-strand break (DSB) formation (Lam and Keeney, 2015). Subsequent steps of DSB repair leads to the formation of reciprocal, interhomologue exchanges known as crossovers (COs)—visualised cytologically as chiasmata—that are essential for faithful disjunction of meiotic chromosomes during anaphase I (Gray and Cohen, 2016; Berchowitz and Copenhaver, 2010). Failure to form at least one CO per homologue pair risks the formation of aneuploid gametes, and thus the process of CO formation is highly regulated (Shinohara et al., 2008; Martini et al., 2006). Importantly, many DSBs—in some organisms the bulk of recombination events—repair without reciprocal exchange and are termed non-crossovers (Storlazzi et al., 1995; Baudat and de Massy, 2007a; Li et al., 2019).

Within many organisms, including *Saccharomyces cerevisiae* (the subject of this study), *Mus musculus*, *Homo sapiens* and *Arabidopsis thaliana*, two subclasses of CO co-exist. ZMM (Zip2-Zip3-Zip4-Spo16, Mlh1-Mlh3, Msh4-Msh5, Mer3)-dependent class I COs account for the majority of COs formed (~70-85% within *S. cerevisiae* (Lynn et al., 2007; de los Santos et al., 2003)). Class I COs are distributed more evenly along each chromosome than expected by chance via a process referred to as CO interference in which the formation of COs in proximity to one another appears suppressed (Berchowitz and Copenhaver, 2010). Class I CO formation requires the nuclease activity of Mlh1-Mlh3, a heterodimer otherwise involved in MMR (Zakharyevich et al., 2012; Cannavo et al., 2020; Kulkarni et al., 2020; Pannafino and Alani, 2021). The less abundant class II COs are non-interfering and depend upon the Mus81-Mms4, Yen1, or Slx1-Slx4 structure-specific nucleases (Holloway et al., 2008; de los Santos et al., 2003; Matos et al., 2011; Sarbajna et al., 2014).

Current models of class I CO formation suggest at least a two-step process involving, initially, the designation of a subset of precursor DSB intermediates by pro-class I CO factors (Bishop and Zickler, 2004). Although CO interference is proposed to become active at this stage, thereby shaping the distribution of COs along chromosomes, the final distribution of class I events is secondarily impacted by a later process: class I CO maturation (Zhang et al., 2014a). Rates of maturation less than 100% will specifically deplete class I COs. Such depletion can lead to achiasmatic chromosomes, or chromosomes with a residual CO positioned close to one or other telomere, where they are considered an “at-risk” location for successful segregation (Wang et al., 2017). Such ideas have been developed to help explain the relatively high rate of meiotic chromosome missegregation in human females relative to males (Wang et al., 2017).

Homologous recombination, the process responsible for both CO and NCO formation, requires a repair template with near-perfect sequence identity (Harfe and Jinks-Robertson, 2000). Mechanistically, Msh2, an essential MMR protein and orthologue of bacterial MutS (Reenan and Kolodner, 1992), is key in binding mismatches to promote their repair through the endonucleolytic action of the Mlh1-Pms1 and Mlh1-Mlh3 complexes (Kolodner and Marsischky, 1999; Srivatsan et al., 2014). Such mismatch detection by Msh2 is thereby integral to preventing recombination by mechanisms that include recruitment of the anti-recombination helicase Sgs1 to heteroduplex DNA (Goldfarb and Alani, 2005). As such, in MMR-proficient *S. cerevisiae* hybrids, recombination between polymorphic homoeologous substrates is inefficient, leading to reduced rates of meiotic CO formation, reduced spore viability, and increased chromosomal non-disjunction during meiosis I (Chambers et al., 1996; Hunter et al., 1996)—phenotypes linked to incipient speciation, and which are largely reversed within MMR-deficient strains (Greig et al., 2003; Hunter et al., 1996; Martini et al., 2011; Chambers et al., 1996; Spies and Fishel, 2015; Bozdag et al., 2021). However, whilst Msh2-dependent binding to mismatches it likely to be rapid (Zhai and Hingorani, 2010), the precise stage of the HR pathway(s) that mismatch recognition arises, and whether this is the same for all intermediates, and in all organisms, is unclear. Indeed, recent observations in mouse meiosis suggest only limited impact of MMR activity on recombination suppression (Peterson et al., 2020), and perhaps more intriguingly, observations in *Arabidopsis thaliana* indicate an unexpected association (rather that inhibition) of recombination within polymorphic regions (Blackwell et al., 2020). Given the fundamental intimate relationships between MMR, sequence divergence, and meiotic recombination, such observations highlight the need to thoroughly explore, and to understand better, the impacts that meiotic MMR activity may have across biology.

## Results

### Deletion of *MSH2* alters the genome-wide frequency and distribution of meiotic crossovers

In order to investigate the impact sequence divergence and the process of mismatch repair has upon meiotic CO formation in the budding yeast, *S. cerevisiae*, we mapped recombination patterns genome-wide (**Fig. 1a** and Methods) within six wild-type and thirteen MMR-defective *msh2*Δ meioses—obtained from a hybrid of two widely utilized laboratory isolates: S288c and SK1 (~65,000 SNPs, ~4,000 high confidence INDELs, ~0.57% divergence; (Martini et al., 2011; Marsolier-Kergoat et al., 2018)). Additionally, we reanalyzed datasets comprising fifty-one wild-type (Mancera et al., 2011; Chen et al., 2008) and four *msh2*Δ (Oke et al., 2014) tetrads from a S96 × YJM789 hybrid of *S. cerevisiae* (~0.6% divergence). On average, we identified 74.3 ±5.4 and 105.9 ±7.8 COs per meiosis within our SK1 × S288c wild-type and *msh2*Δ samples respectively, corresponding to a significant 1.4-fold increase in CO frequency (p<0.01; Two-sample T-test) (**Fig. 1b** and **Supplementary Table 1**). A significant *msh2*Δ-dependent increase (~1.25-fold, p<0.01; Two-sample T-test) was also observed within the S96 × YJM789 hybrid (**Fig. 1b** and **Supplementary Table 1**)—collectively reaffirming the known anti-recombinogenic activity of Msh2. Notably, CO frequencies were considerably higher within S96 × YJM789 than S288c × SK1 (91.4 vs. 74.3 COs per wild type meiosis)—suggesting that cross-specific differences may exist.

**Fig. 1.**
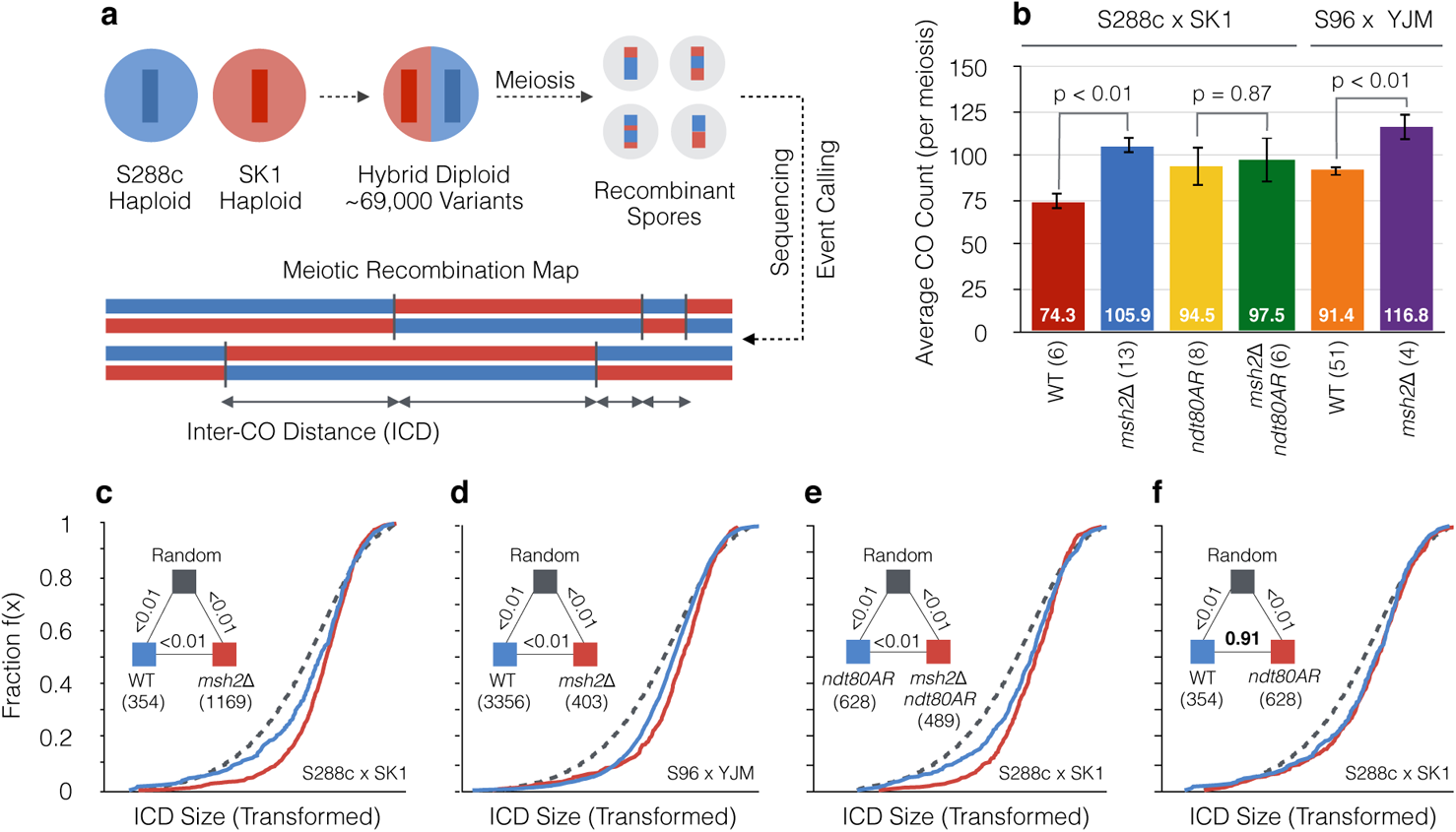
Inactivation of Msh2 increases CO frequency and the global strength of CO interference. **a,** Genome-wide mapping of recombination. Meiosis is induced within hybrid S288c × SK1 *S. cerevisiae* diploid cells and genomic material is prepped from individual, isolated spores for paired-end Illumina sequencing in order to genotype SNP/indel patterns and therefore determine the parental origin of any given loci (**Methods**). Only a single chromosome is shown for clarity. Inter-crossover distances (ICDs), a measure of the uniformity of CO distribution, are calculated as the distance (in bp) between successive COs along a given chromosome. **b,** Average number of COs per meioses for each genotype. The number of individual meioses sequenced per genotype is indicated. Error bars: 95% confidence intervals (CI). P values: Two-sample T-test. **c-f,** Empirical cumulative distribution functions (eCDFs) showing the fraction of ICDs at or below a given size. The total number of experimental ICDs is indicated in brackets. ICDs are transformed (**Methods**) to correct for skews generated by differing CO frequencies. Untransformed ICD plots are available in (**Supplementary Fig. 1c-h**). Randomised datasets were generated via simulation to represent a state of no interference (Methods). Pairwise goodness-of-fit tests were performed between genotypes as indicated (triangular legend). P values: Two-sample KS-test.

Deletion of *MSH2* also increased observed NCO events in both hybrid crosses (SK1 × S288c: wild type = 30.8; *msh2*Δ = 92.8; S96 × YJM789: wild type = 46.8; *msh2*Δ = 56.3; **Supplementary Table 1**), supporting the established effect of Msh2 in suppressing both COs and NCOs in hybrid strains as reported previously (Martini et al., 2011). However, unlike COs, the visibility of NCOs is directly affected both by the true number of converted and/or heteroduplex markers contained within a NCO event, and by the technical efficiency of calling what may potentially be only short regions of contiguous nonreciprocal marker change (Marsolier-Kergoat et al., 2018; Ahuja et al., 2021). NCO frequencies are also more affected by homeostatic effects than are COs (Martini et al., 2006). For these reasons, quantitative comparisons of NCO frequency changes in the presence and absence of *MSH2* are not simple to interpret. Instead, we focused our attention on COs where event visibility is assumed to be similar in the presence and absence of Msh2.

To investigate any possible effect of Msh2 on CO patterning, we determined the distribution of inter-crossover distances (ICDs)—the separation (in bp) between successive COs along every chromosome. To accommodate comparisons between different sample sizes, ICDs were transformed (**Methods** and **Supplementary Fig. 1**) and visualized in rank order as empirical cumulative distribution functions (eCDFs). In all mapped strains, CO distributions deviated significantly (p<0.01; Two-sample Kolmogorov-Smirnov (KS) test (Massey Jr, 1951)) from simulated conditions in which the same frequency of observed COs were distributed randomly relative to one another (**Fig. 1c–f**). In particular, observed patterns displayed a more homogenous distribution of ICDs, with fewer short and long distances between adjacent COs than expected by chance. This change in distribution was visible as a steeper eCDF curve in the experimental data compared to the random simulation, a feature that was stronger within the S96 × YJM789 cross (**Fig. 1d**) than within S288c × SK1 (**Fig. 1c**), additionally suggesting that the meiotic CO landscape is regulated in a cross-specific manner (**Supplementary Fig. 2** and **Extended discussion**).

Unexpectedly, deletion of *MSH2* within both hybrid crosses caused ICD curves to skew yet further away from the random simulation, creating a steeper inflection point (**Fig. 1c-d**; p < 0.01; Two-sample KS test) indicative of increased homogeneity in the spacing of COs relative to one another. Removal of a second MMR factor, *PMS1*, which acts downstream of Msh2 mismatch binding (Prolla et al., 1994), caused similar changes to both the CO frequency and the CO distribution curves relative to wild type (**Supplementary Fig. 3a-b**), but no additive effect in a double *msh2*Δ *pms1*Δ strain was observed—thereby suggesting that these CO changes arise as a general consequence of MMR inactivation.

To account for any impact increased CO frequency may have upon CO distribution, we utilized an *ndt80AR* (“arrest–release”) strain where meiotic prophase length is extended via temporary repression of the Ndt80 transcription factor (Benjamin et al., 2003; Xu et al., 1995; Crawford et al., 2018). On average, we identified 94.5 ±16.3 COs per meiosis within *ndt80AR*—a significant increase relative to wild type (p<0.01; Two-sample T-test) (**Fig. 1b**). However, no further increase occurred upon deletion of *MSH2* (*msh2*Δ *ndt80AR*, 97.5 ±15.4 COs, p=0.87; Two-sample T-test) (**Fig. 1b**). Importantly, despite the lack of change in CO frequency, the *msh2*Δ-dependent skew in CO distribution was still observed in *ndt80AR msh2*Δ (**Fig. 1e**) whereas increased CO frequency alone (*ndt80AR*) did not alter CO distribution compared to wild type (**Fig. 1f**) (p=0.91; Two-sample KS test).

Inactivation of MMR within hybrid *S. cerevisiae* strains therefore gives rise to two distinct phenotypes relative to wild type: (i) increased CO frequency, as previously observed (Martini et al., 2011), and (ii) a global shift in the distribution of COs relative to one another—something that can arise independently of changes in CO frequency. Whilst we are not directly measuring interference between COs by these analyses, the latter change to the CO distribution is consistent with the loss of MMR activity appearing to increase the presence and/or impact of CO interference within the global pool of COs.

### Mixture modelling of class I and class II CO distributions

CO distributions can be modelled by the gamma (γ) distribution, where γ(α) values >1.0 indicate increasing deviations from randomness (potentially indicating increasing strength of interference between COs) (McPeek and Speed, 1995). Importantly, however, because the experimentally observed CO distribution is a composite mixture of class I (interfering) and class II (non-interfering) COs, models based on a single (γ) distribution deviate substantially from experimental data, which contains many more ICDs <50 kb than expected (**Fig. 2a**). Indeed, the presence of these short ICDs is consistent with the presence of the subpopulation of randomly distributed COs (i.e. class II).

**Fig. 2.**
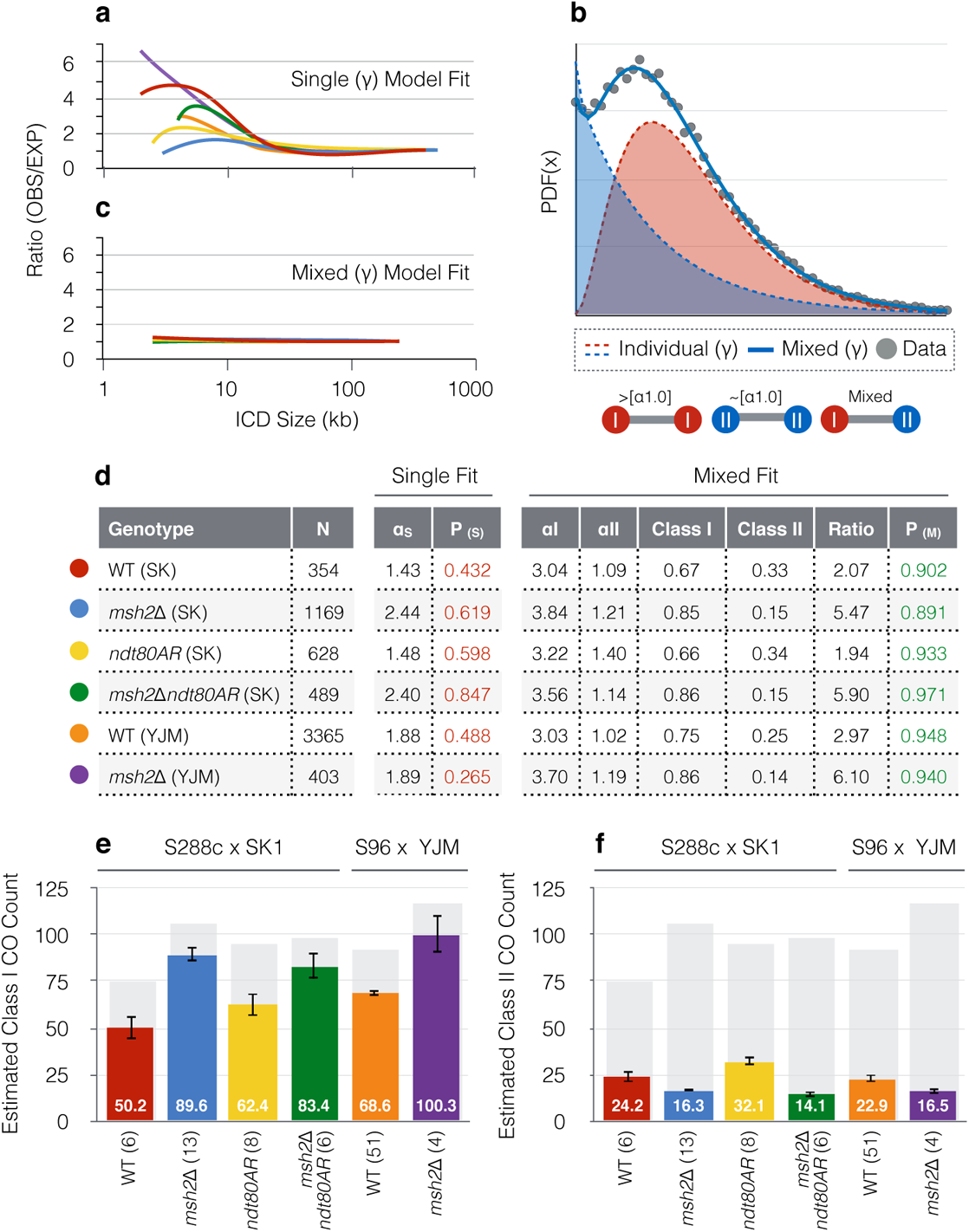
Computational modelling predicts a Msh2-dependent shift in the class I:class II CO ratio. **a,** Ratio of experimentally observed ICD sizes (OBS) versus the theoretical expectation based on a single, best-fit gamma (γ)-distribution (EXP). Ratio values were calculated at 5 kb intervals. **b,** Example (γ) mixture model (α1.0|β1.0 + α3.0|β 5.0). I = class I. II = class II. k datasets, owing to the existence of two CO subclasses, are a heterogenous population of three ICD types (as shown). **c,** As in **(a)** but based on a mixed (γ)-model (no. of distributions fitted = 2). **d,** Best-fit (γ) mixture modelling results. N = sample size (total number of ICDs). α_s_ = Single-fit γ(α) value. P_(s)_ = Fit quality of a single (γ)-distribution (one-sample KS-test). αI, αII = Mixed model γ(α) values for each class. Class I, ClassII = estimated fraction of each CO subclass. Ratio = class I:class II. P_(M)_ = Fit quality of a mixed (γ)-mode (Two-sample KS-test) **e-f,** Estimated class I and class II CO counts respectively. Estimates were obtained using the best-fit class I:class II ratios. Total CO frequencies are overlaid (grey bar). Error bars: 95% confidence intervals. The number of individual meioses sequenced per genotype is indicated.

To explore this aspect of the data, we utilised a computational method based on (γ)-mixture modelling (**Fig. 2b**), to statistically deconvolute ICD data, thereby deriving estimates of the random and non-random components (**Methods**). A similar approach has been developed by others to explore the parameters governing the physical positioning of COs along chromosomes (Gauthier et al., 2011). A non-random (γ(α_I_) >3.0), and a random (γ(α_II_) ~1.0) component was identified for all genotypes (**Fig. 2d**). Composite simulations (**Supplementary Fig. 4**) using the estimated proportions of these two CO distributions improved model fit to experimental data and eliminated the deviations between simulation and experimental data otherwise observed for ICDs below 50 kb (**Fig. 2c**). We henceforth refer to these random and non-random CO components as estimates of, respectively, the class II and class I CO components.

Consistent with prior estimates (de los Santos et al., 2003), using this analysis, *MSH2* wild type class I:class II ratios were estimated at ~2.0 and ~3.0 within S288c × SK1 and S96 × YJM789 respectively (**Fig. 2d**). By contrast, deletion of *MSH2* increased ratio estimates to ~5-6 in both hybrid crosses (**Fig. 2d**), suggesting a ~1.7-fold increase in class I CO formation in the absence of Msh2 (**Fig. 2e**). These models also estimated *msh2*Δ-dependent decreases in the absolute frequency of class II CO formation (~0.5 to 0.7-fold) (**Fig. 2f**). Importantly, the γ(α_I_) value estimates obtained for the non-random (class I) CO population were broadly similar for all strains, irrespective of Msh2 status or estimate of total class I CO frequency (gamma values between ~3.0–3.8; **Fig. 2d**, Mixed fit)—and different from the poorly fitting global γ(α_s_) estimates (1.43–2.44) generated from fitting a single gamma distribution to each entire dataset (**Fig. 2d**, Single fit).

Thus, assuming that our modelling of a mixed gamma distribution is a reasonable estimate of the class I : class II balance present within cells, these analyses suggest that the global change in CO distribution towards less randomness in *msh2*Δ arises not from an increase in CO interference strength between class I COs, but instead from a change in the relative proportion of class I and class II COs present within the total CO pool. Specifically, our results suggest that the formation of ZMM-dependent class I COs is preferentially favoured within Msh2-deficient cells and that most of the additional COs observed within *msh2*Δ relative to wild type are class I. Put another way, our observations suggest that the activity of Msh2 disproportionately impedes the formation of class I COs relative to class II.

### Distribution and frequency of Zip3 foci appear unaltered in *msh2*Δ

Zip3 foci specifically mark the sites of interfering, class I COs (Agarwal and Roeder, 2000; Zhang et al., 2014b). Our computational analysis predicts an increase in Zip3-marked CO sites, but no change in their relative distribution along synapsed chromosome axes. Thus, to investigate the results of our modelling, we counted Zip3 foci (Shinohara et al., 2008) on spread meiotic chromosome preparations from S288c × SK1 hybrids co-labelled with Zip1-GFP (White et al., 2004), an established marker of chromosome synapsis in *S. cerevisiae* (**Fig. 3a-b** and **Supplementary Fig 5**) (Henderson and Keeney, 2004). To reduce observational bias, samples were randomised, and counting was restricted to only well-spread nuclei showing clear thread-like patterns of Zip1. Overall, focus number per nucleus was highly variable (10–50 per cell). Perhaps because of this variation, we were unable to identify any reproducible differences in foci number upon *MSH2* deletion, both in the wild-type and the pachytene-arrested *ndt80AR* strain background. However, we cannot exclude that this high variance obscures a real difference.

**Fig. 3.**
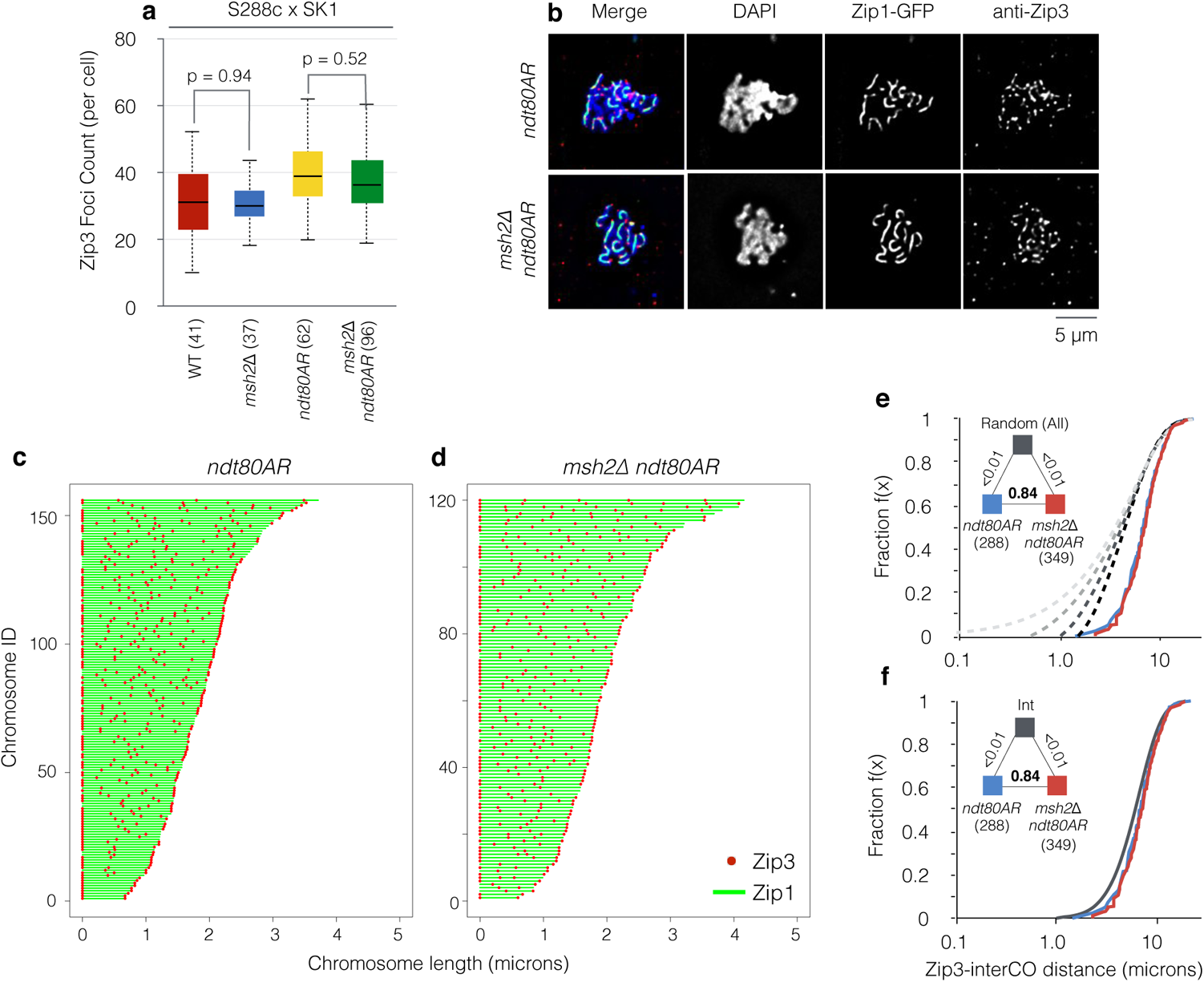
Zip3 foci counts are neither elevated nor redistributed within Msh2-deficient cells. **a,** Box-and-whisker plot showing Zip3 foci counts obtained from chromosome spreads of S288c × SK1 *ndt80AR* cells prepared at 8 h following induction of meiosis (pachytene arrest). Midlines denote median values, box limits are first and third quartile, whiskers are highest/lowest values within 1.5-fold of interquartile range. P values: Two-sample T-test. **b,** Representative example for each genotype. Cells are fluorescently labelled for the meiosis-specific axis protein Zip1-GFP (green), the class I CO marker, Zip3 (red) and DNA (DAPI, blue). Only well-spread nuclei with clear Zip1 threads were analysed. Only Zip3 foci overlapping within the DAPI-stained area were counted. The total number of nuclei counted is indicated in brackets obtained from three independent experiments. **c-d,** Relative distribution of Zip3 foci along individual Zip1-GFP positive chromosome axes in *ndt80AR* and *msh2*Δ *ndt80AR* pachytene-arrested cells, ordered from bottom to top by increasing axis length (Green bar, measured Zip1 axis length; red dot Zip3 focus position). **e-f,** Inter-Zip3 foci distances (measured in microns) were aggregated, rank ordered, and expressed as a fraction of the total (i.e. an eCDF), equivalent to our presentation of inter-CO distances detected from octad sequencing data. In (**e**), observed distributions are compared to simulations of the same number of randomly distributed foci over the same total axial distance, using four increasingly stringent merging thresholds (0.05, 0.01, 0.15, and 0.2 microns; light grey to dark grey dashed lines) equivalent to approximate range of imaging resolution (1 pixel = 0.1 microns). Observed distributions were not statistically dissimilar in the presence and absence of Msh2, but were significantly different from all simulated random distributions regardless of merging threshold. In (**f**), the observed distributions are compared to a simulated gamma distribution, that whilst still statistically dissimilar, shows a clear visual similarity. The residual deviation from an interfering gamma distribution may be caused by inherent inaccuracies in microscopy resolution, or a real characteristic of Zip3 foci as measured along spread chromosome axes.

Within these data we noticed a modest correlation between spread size and total foci count per spread (**Supplementary Fig. 6a**). Thus, to investigate whether technical differences in spreading efficiency was affecting Zip3 foci counts we further analysed the density of Zip3 foci per µm^2^ of spread area bounded by the chromosomal (DAPI) signal, but, again, found no clear differences caused by *MSH2* deletion (**Supplementary Fig. 6b**).

We next analysed the relative distribution of Zip3 foci along synapsed chromosome axes—a potential indicator of CO interference—in cells arrested at the pachytene stage via the *ndt80AR* allele (**Fig. 3c-f**). Measuring the distances between Zip3 foci along the subset of well-resolved chromosomes in each strain demonstrated significant deviation from that expected for a random distribution (**Fig. 3e**) and instead much closer to that of a simulated interfering distribution (**Fig 3f**), as expected for Zip3 foci marking interfering Class I events (Zhang et al., 2014b). However, no additional distributional differences were detected in the absence of Msh2 (**Fig. 3e-f**). The similarity of the Zip3 foci distribution in the presence and absence of Msh2 is consistent with our mixture-modelling analysis of CO positions identified in the genome-wide data, which estimated a similar γ(α_I_) parameter for the non-random component in all strains regardless of *MSH2* status (**Fig 2d**, above).

Whilst the large variation in Zip3 foci frequencies limits our ability to draw firm conclusions, taken together, our observations suggest that the differences in CO frequency and patterning that are observed in our genome-wide analysis with and without *MSH2* may arise downstream of the point at which we have assessed Zip3 foci counts (“pachytene” as mediated by *ndt80*Δ arrest). Specifically, changes in the efficiency of CO designation and/or maturation arising after this arrest point, and/or independent of Zip3 focus formation, could lead to differences in observed (Zip3 foci) and final (genome-wide analysis in tetrads) CO frequency and distribution. This possibility is explored in more detail below.

### Msh2 specifically impedes class I CO formation at regions of greater sequence divergence

The MMR machinery forms a potent barrier to homoeologous recombination, presumably due to recognition and destabilisation of recombination intermediates containing DNA mismatches. To directly explore the interplay between DNA mismatches and CO formation, we calculated polymorphism densities (SNPs, Indels) ±500 bp around every mapped CO (**Fig. 4a**) and compared between genotypes. To generate a comparative reference point, the expected environment for meiotic recombination, as defined by the polymorphism density surrounding ~3600 DSB hotspot midpoints (Pan et al., 2011), was also calculated. Polymorphism density surrounding COs within *MSH2* wild-type strains (S288c × SK1: wild type, *ndt80AR*) was significantly different to that expected in both distribution (p<0.01; Two-sample KS test) and in mean variant density (5.32 vs. 6.18, p<0.01; Two-sample T-test)—characterized by a skew towards COs, on average, arising within regions of lower genetic divergence than expected from the genome-wide position of DSBs (**Fig. 4b**). By contrast, the polymorphism density around *msh2*Δ COs displayed visual and statistical similarity to that expected (p>0.25; Two-sample KS-test) (6.26 vs. 6.18, p = 0.52; Two-sample T-test) (**Fig. 4b**). Such a disparity between wild type and *msh2*Δ was recaptured within the independent S96 × YJM789 hybrid where COs were again skewed towards regions of lower polymorphism density only in the *MSH2* wild-type strain (**Fig. 4c**). A similar effect was observed when considering polymorphism density arising ±1000 bp around each CO but was diminished with increasing distance (±2000 bp)—suggesting that DNA mismatches exert a localised inhibitory effect on CO formation (**Supplementary Fig. 7a-b** and **Supplementary Discussion**).

**Fig. 4.**
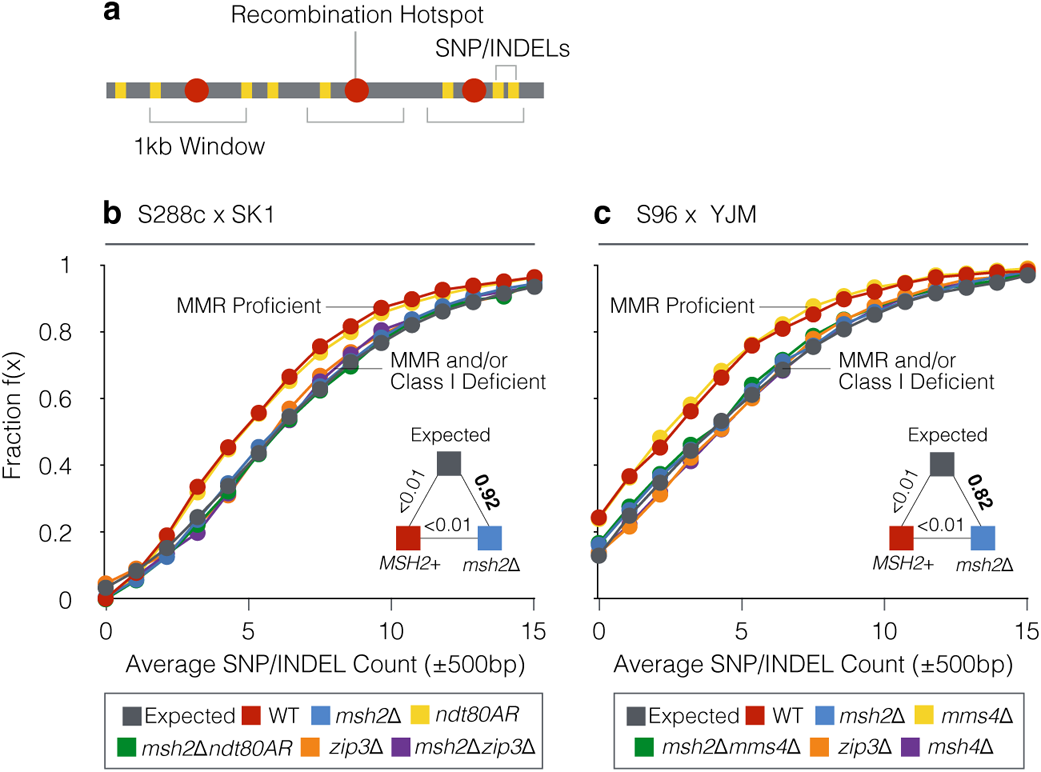
Suppression of class I COs occurs at regions of higher sequence divergence. **a,** SNP/indel count is assayed using a ±500 bp window centred on CO or DSB hotspot midpoints. All contained SNP/indels are tallied with equal weight. **b-c,** Empirical cumulative distribution functions (eCDFs) showing the fraction of COs that reside within a region of a given SNP/indel count for S288c × SK1 (**b**) and S96 × YJM789 (**c**) for the indicated hybrid strains. Expected eCDF curve (grey) is calculated using DSB hotspot midpoints (Pan et al 2011). Pairwise goodness-of-fit tests were performed between pooled *msh2*Δ and *MSH2* control datasets as indicated (triangular legend). P values: Two-sample KS-test.

Our mixture modelling suggests that inactivation of *MSH2* alters global CO distribution by altering the relative abundance of class I vs class II COs (**Fig. 2** and **Fig. 3**). To determine whether the influence that Msh2 has in regulating CO class outcome is related to its role in suppressing COs at sites where mismatches will arise during meiotic recombination, we further calculated polymorphism densities within mutants that disrupt class I (*zip3*Δ, *msh4*Δ) or class II (*mms4*Δ) CO formation (Oke et al., 2014). Strikingly, mutants devoid of class I COs phenocopied *msh2*Δ—that is, COs within these mutants were no longer skewed away from regions of higher polymorphism density despite the presence of Msh2—sharing mean polymorphism densities around COs that were not statistically dissimilar to expected (p>0.5; Two-sample T-test) (**Fig. 4b-c**). Moreover, the impact of *zip3*Δ and *msh2*Δ appeared to be epistatic rather than additive, with no further change in the double mutant (**Fig. 4b**), suggesting a single common pathway. By contrast, removal of class II formation (*mms4*Δ) had no impact on the interplay between CO formation and polymorphism density (**Fig. 4c**)—collectively suggesting that mismatch-dependent repression of CO formation is specific to class I COs.

### Msh2 activity during class I CO maturation

Whilst our observations provide strong evidence to support a disproportionate impact of Msh2 and polymorphisms on COs formed via the class I pathway, the time and mode of action within the HR process that this occurs is unclear. To explore in more detail the possibility of early (designation) versus late (maturation) action of Msh2, we generated simulations in which mixed patterns of COs (ranging from 100% class I to 50% class I : 50% class II) were subject to varying rates of stochastic class I CO failure (up to 50% loss)—simplistically mimicking the possible outcome of class I CO maturation failure at a (late) stage downstream of interference-patterned CO designation (**Fig. 5a** and **Methods**). Resulting genome-wide patterns of simulated CO formation (presented as eCDF curves of inter-CO distances as in **Fig. 1c-f**) were then compared to our experimentally observed CO patterns and tested for statistical similarity (**Fig. 5b-g**). For *msh2*Δ datasets, in both SK1xS288c (**Fig. 5b-c**) and S96xYJM (**Fig. 5d**), better fits (P>0.9, KS test) were obtained only for parameter combinations in which either class II CO formation was low (as described above) and/or maturation failure was low—consistent with the relatively regular (interfering) genome-wide pattern of COs observed in these strains (**Fig. 1c-f**).

**Fig. 5.**
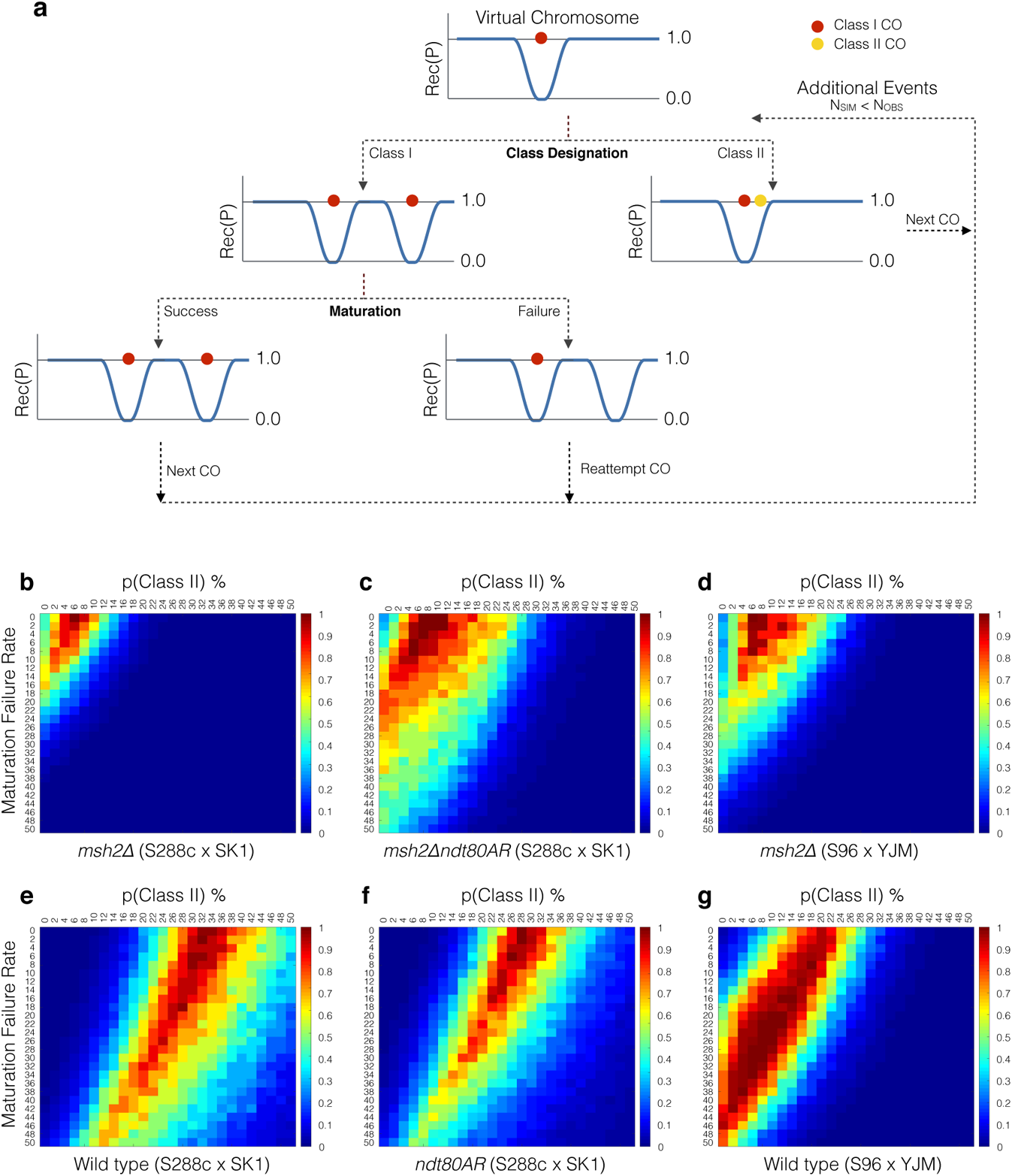
Simulating impact of CO maturation failure on observed CO distributions. **a,** Extended *RecombineSim* platform as described in **Supplementary Fig. 4**, but with the introduction of variable rates of stochastic class I CO maturation failure downstream of CO interference patterning in addition to variable fractions of randomly distributed class II COs. In this simulation, class I COs that fail to mature are still sensitive to, and still generate, localised regions of interference, but are removed from the final observed pattern of visible events. In such instances, additional COs are simulated until the final observed simulated frequency matches the frequency observed in experimental datasets. COs arising within 1.5 kb of one another are merged into a single event, again matching the way experimental datasets are processed. **b-g,** Coloured heat maps of P values (KS test) between observed and simulated CO distributions expressed as eCDF curves for the indicated strains. P values >0.9 indicate good statistical fits. Each pixel represents a particular combination of parameter values: maturation failure rate (Y axis) and class II CO % (X axis). See main text and Methods for more details.

By contrast, and as described above, wild-type cells, where Msh2 is active, displayed global CO patterns that were better fit (P >0.9; KS Test) with a greater proportion of class II COs (**Fig. 5e-g**; ~30% in SK1xS288c; ~20% in S96xYJM). However, these simulations also identified a diagonal band of reasonable parameter fits (P >0.8; KS Test) where decreasing proportions of class II COs were offset by increasing chances of class I CO maturation failure (**Fig. 5e-g**). This trend was clear, albeit relatively modest, in the SK1xS288c hybrid, but was enhanced in S96xYJM, where the highest density of good statistical fits (P>0.9; KS test) was obtained for relatively low fractions of class II COs (~5-10%), similar to the estimates obtained in *msh2*Δ cells, but with high rates of class I CO maturation failure layered on top (20-40%; **Fig. 5g**).

Collectively, our simulations suggest that intrinsic rates of class I maturation failure may be quite low in *msh2*Δ hybrid cells. By contrast, mismatch-dependent class I maturation failure, downstream of CO designation, and thus potentially quite late in meiotic prophase, may underpin at least some of the apparent reductions in class I COs observed in *MSH2* control cells—an effect that may be more prevalent in certain hybrids such as S96xYJM.

## Discussion

Sequence divergence suppresses recombination within a wide range of eukaryotes including *S. cerevisiae*, *M. musculus* and *H. sapiens* (Chambers et al., 1996; Hunter et al., 1996; Bozdag et al., 2021; Cole et al., 2010; Baudat and de Massy, 2007b; Jeffreys and Neumann, 2005). Findings presented here expand upon these observations and suggest that the anti-recombinogenic activity of Msh2, exerted at homoeologous sites, does not mediate an indiscriminate suppression of COs but rather acts disproportionately at sites of ZMM-dependent, interfering COs (**Fig. 6a-c**)—thereby altering the spatial distribution of CO recombination across the genome by modulating the class I : class II balance. Our observations underscore how even the low rates of divergence (~0.6%) present within intra-specific hybrids of *S. cerevisiae* can generate global changes in CO frequency, CO type, and genome-wide distribution. Nevertheless, in wild-type hybrids when Msh2 is active, COs still frequently arise within heterologous regions (**Fig. 4b-c**) with class I COs still forming at a high rate (67%-75% of total COs; **Fig. 2d**). Mismatches do not, therefore, form an absolute barrier to class I COs, but instead seem to influence the probability of their formation.

**Fig. 6.**
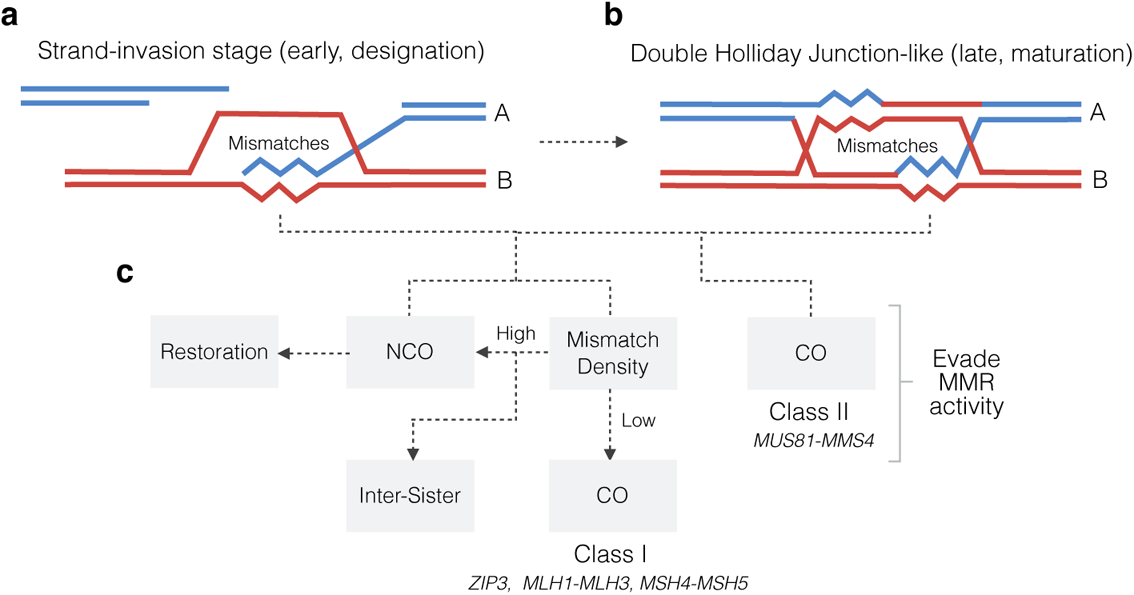
Model summarising mismatch-directed suppression of class I COs. **a-b,** Mismatches (jagged lines) may arise within recombination intermediates at various stages of the meiotic recombination pathway due to differences in sequence between parental information A (blue) and B (red). **c,** In the presence of a functional MMR pathway, regions of higher sequence divergence are proposed to give rise to transient heteroduplexes that cause Msh2-dependent redirection of repair toward the NCO or inter-sister outcomes. Formation of NCOs could arise via destabilisation of nascent strand-invasion intermediates, dissolution of dHJs via Sgs1-Rmi1-Top3, or disruption of CO-biased dHJ resolution (see text for more details). Concomitant repair of mismatches may additionally render some NCO events invisible, and thereby indistinguishable from inter-sister events, due to restoration of parental markers. Inactivation of MMR alleviates such repression arising within CO precursors, increasing the frequency of class I CO formation and thus the spatial uniformity of relative CO positions. In the absence of pro-class I CO factors such as Zip3, Mlh1-3 and Msh4-5, class II COs can arise, but are less subject to Msh2-dependent destabilisation, perhaps due to intrinsic differences in structure, lifespan, and/or extent of heteroduplex DNA. For example, extended branch migration of Holliday junctions at class I precursors (Marsolier-Kergoat et al 2018) which may stabilise such intermediates (Ahuja et al 2021) could increase the probability of hDNA arising within them, and thereby increase Msh2-dependent redirection towards NCO outcomes.

Current models posit that the establishment and differentiation of class I COs is a multi-step process starting with nascent recombination interactions initiated by Spo11 DSBs, designation as class I precursors, and subsequent maturation into the final class I COs (Zhang et al., 2014a). Inefficient maturation—downstream of the implementation and patterning effects of CO interference—has been proposed as a mechanism to explain CO patterns and the innate predisposition towards meiotic chromosome missegregation in human females (Wang et al., 2017).

Within this framework, it is therefore important to consider at what stage Msh2 activity causes class I CO precursors to be redirected towards alternative outcomes. Prior analyses aimed at elucidating the CO differentiation process have harnessed relatively deep (rather than broad) datasets in order to build probability distributions of expected coincident class I COs at adjacent positions along specific chromosomes (Zhang et al., 2014a). By contrast, our genome-wide maps of CO position are broad, encompassing positional information for every CO on every chromosome in individual meioses, but are of limited depth at any given locus due to the limited throughput of genome-wide sequencing of meiotic progeny (six wild type and thirteen *msh2*Δ meioses in the S288c × SK1 hybrid). These differences preclude us from performing a similar analysis. Moreover, unlike the specificity of Zip3-focus analysis for class I COs, genome-wide CO maps cannot yet distinguish between class I and class II COs at any given site. For these reasons, we have focused on analysing Msh2-dependent differences in the fine-scale positions of COs and in the distribution of COs relative to one another.

When considering global CO positions, conversion from a class I to a class II CO would still give rise to a CO in the same position and with no change in global CO frequency. Thus, because we observe changes in CO pattern and frequency, we infer that most Msh2-redirected class I events become NCOs and/or become otherwise invisible within our assay (**Fig. 6c**). On this latter point, recent work provides evidence for frequent repair-template switches between homologues and sister chromatids (Marsolier-Kergoat et al., 2018; Ahuja et al., 2021), and thus redirection of CO precursors towards repair exclusively using identical sequences on the sister chromatid is also possible (events that would be invisible in our assays). However, such redirection would seemingly need to happen prior to the priming of DNA synthesis by a DSB end that has invaded the homologue (**Fig. 6a**). Alternatively, Msh2-dependent redirection towards inter-sister events and NCOs may occur concomitantly with mismatch repair, leading in some cases to restoration of any heteroduplex markers back to the parental configuration, again precluding detection by our methods (**Fig. 6c**).

The fact that the estimated frequency of class II COs was greater, in absolute terms, in Msh2-proficient cells suggests that Msh2-dependent suppression of class I COs may indirectly influence class II CO formation. For example, more class II events (arising from increased Spo11 activity) might be enabled by spatial and/or temporal changes in the efficiency of homologue engagement caused by MMR-dependent rejection of nascent recombination events, similar to when class I CO formation (Thacker et al., 2014) and/or chromosome synapsis (Mu et al., 2020) is disturbed by genetic mutation. However, we did not detect any obvious Msh2-dependent differences in synapsis within hybrid strains (**Fig. 3b**), suggesting that, if present, such effects are transient and/or relatively subtle.

In our envisioned model (**Fig. 6a-c**), which builds upon prior ideas (Chambers et al., 1996; Hunter et al., 1996), rejected class I events may be redirected towards the NCO pathway relatively early on during CO designation and/or maturation via unwinding of the initial strand invasion intermediate with repair proceeding via synthesis-dependent strand annealing (SDSA; (Marsolier-Kergoat et al., 2018; Ahuja et al., 2021)). Alternatively, redirection could occur at a later stage, for example after class I CO precursors reach the double Holliday junction (dHJ) stage. The potential for dHJs to undergo branch migration (Marsolier-Kergoat et al., 2018; Ahuja et al., 2021) may generate large patches of heteroduplex DNA that could in turn be efficiently detected by the MMR machinery and stimulate Sgs1-Top3-Rmi1-dependent dissolution. It is also possible that mismatches cause dHJs to be more-frequently resolved as NCOs nucleolytically (perhaps via nicks generated by the MMR process itself), whereas an almost absolute bias towards CO resolution is observed in the absence of Msh2 (Marsolier-Kergoat et al., 2018).

Our apparent inability to detect Msh2-dependent differences in the frequency of Zip3-marked CO precursors at the pachytene-like arrest enforced by *NDT80* deletion where dHJs accumulate (**Fig. 3**; (Allers and Lichten, 2001)), indeed suggests that at least some class I redirection may arise after this dHJ stage. A late-stage activity of Msh2 that disturbs class I CO formation would also be compatible with our simulations of CO maturation failure in wild-type, but not *msh2*Δ cells (**Fig. 5**). However, why the two hybrids studied here behaved differently in this regard is unclear but might suggest differing propensity for Msh2 to elicit anti-CO effects before, during, or after CO patterning has completed.

It is also possible that the distributional differences in CO patterns we have observed are patterned by processes that are independent of CO interference and the class I or class II CO pathways. Non-uniform densities of DNA-sequence polymorphisms, DSBs, and even COs themselves all have the potential to influence the relative CO distributions that arise on a per-cell basis. However, polymorphisms, DSBs, and COs are relatively evenly spread across the entire length of each chromosome in *S. cerevisiae* (**Supplementary Fig. 8a-d**), and thus as expected, biasing CO site selection by these underlying population-level parameters had no measurable impact on resulting patterns of simulated inter-CO distributions (**Supplementary Fig. 8e-g**). Nevertheless, we recognise that non-random distributions of precursor events in individual cells can influence downstream patterns of COs (Zhang et al., 2014a), without necessarily generating nonuniformity when assayed across a population. In addition, in organisms with less uniform SNP/indel density, and/or propensity to initiate recombination, such non-uniformity could result in a redistribution of CO formation towards certain regions, potentially influencing relative CO patterning on a per-cell basis (e.g. the effect that highly heterologous regions have in *A. thaliana* (Ziolkowski et al., 2015)).

In mitotic cells, inhibition of homoeologous recombination by means of heteroduplex rejection, relies upon Msh2 and the RecQ-family helicase, Sgs1 (Sugawara et al., 2004; Goldfarb and Alani, 2005; Spies and Fishel, 2015). An *sgs1*Δ mutant may therefore be expected to phenocopy *msh2*Δ if suppression of class I COs occurs via this mechanism. Intriguingly, however, the distribution of COs is more random in *sgs1*Δ relative to wild type (**Supplementary Fig. 3c**), suggesting a decrease in the proportion of class I COs. Moreover, *sgs1*Δ also abolishes the increased skew towards nonrandomness (inferred above to indicate an increased frequency of class I COs) caused by *MSH2* deletion (**Supplementary Fig. 3d**)—suggesting that Msh2 and Sgs1 are not epistatic, but rather antagonistic in the formation of class I COs. Thus, it is possible that Msh2 mediates suppression of class I COs in a pathway different to that of Sgs1-mediated heteroduplex rejection, instead relying upon the downstream properties or factors of MMR, including Pms1 (**Supplementary Fig. 3a**), to achieve its effect. Alternatively, the genetic complexity outlined above may arise because Sgs1 can act at multiple steps and on a range of recombination intermediates. For example, at early stages Sgs1 could act to promote class I CO formation by unwinding nascent recombination intermediates—independently of Msh2 and mismatches—thereby allowing them to be recycled into future potential class I precursors (Oh et al., 2007; Tang et al., 2015; Kaur et al., 2015). By contrast, perhaps mediated by mismatch- and Msh2-dependent destabilisation of pro-class I CO factors, Sgs1 activity at a later stage could promote dHJ dissolution thereby suppressing class I maturation.

It is important to consider how MMR-specificity for class I COs may arise. Msh2, Mlh1 and Pms1 form a ternary complex during MMR (Li, 2008) and *in vitro* data suggest that Mlh1-Mlh3—essential class I CO factors— facilitate binding of Msh2 to heteroduplex DNA arising, for example, at sites where mismatches exist between parental strains (Rogacheva et al., 2014). Msh2 itself, via interaction with Msh6, also directly binds Holliday junctions with high affinity (Alani et al., 1997). Thus, Mlh1-Mlh3, or the inherent structure of class I precursors, may be responsible for the differential sensitivity of each CO subclass to sequence mismatches through preferential recruitment or activation of Msh2 and MMR at class I sites. Indeed, available data (Getz et al., 2008) suggest that Class I COs are more likely to recruit MMR and lead to conversion/restoration (6:2, 4:4 patterns)—or possibly rejection (our results). By contrast, class II COs are less likely to recruit MMR, and thus not only survive in MMR-proficient cells but also show signs of post-meiotic segregation (Getz et al., 2008). Such mechanisms would fit with the specific reductions in class I COs that we observe in MMR-proficient cells.

Given the evolutionary conservation of MMR and of the fundamental process of CO recombination in meiosis, an important consideration is whether the processes uncovered by our study are conserved across biology. With this in mind, it is interesting to note that a detailed analysis of two Spo11-DSB hotspots in *M. musculus* found no evidence for MSH2-dependent suppression of recombination (Peterson et al., 2020). Furthermore, a recent genome-wide study in *A. thaliana* indicates redistribution of COs towards, rather than away from, polymorphic regions in MMR-proficient control lines relative to a *msh2* mutant (Blackwell et al., 2020). Thus, it will be critical to elucidate what are the mechanistic relationships that underpin these species-specific observations. Such differences may be directed by species-specific modulation of MMR activity at pro-CO sites, or by fundamental genome-scale differences that influence meiotic recombination, such as chromosome size, global recombination rate (high in yeast), distribution of DNA sequence heterozygosity (non-uniform in *A. thaliana*), and regulation of recombination initiation (PRDM9 dependent in *M. musculus*).

Overall, understanding the molecular mechanisms that contribute to speciation is fundamental to our understanding of biological diversity and evolution. In this regard, despite many unknowns remaining, our observations highlight an unexpected link between DNA sequence divergence, MMR, and meiotic class I CO control, thereby intimately linking the regulation of CO numbers and their distribution to pathways contributing to reproductive isolation and eventual speciation.

## Contributions

T.J.C, M.R.C. and M.J.N. conceived of the project. M.R.C. performed all wet-lab work, data processing and event calling associated with the genome-wide mapping. T.J.C. analysed and interpreted the data, performed in-silico simulations, and designed the modelling algorithms. L.J.H. performed all microscopy and foci analysis. M.M.K. and B.L. provided scripts, protocols, additional samples, and ideas. T.J.C., M.J.N., and B.L. wrote the manuscript with critical input from all authors.

## Acknowledgements

We thank A. Shinohara for sharing anti-Zip3 antibody (Shinohara et al., 2008) and R. M. Allison for help with image acquisition.

## Corresponding authors

Correspondence to: Matthew J. Neale and Bertrand Llorente

## Data Availability

Raw sequence data is deposited in the NIH Sequence Read Archive (SRA) under accession numbers <u>SRP151982</u> (wild type, *msh2*Δ, *ndt80AR*), <u>SRP111430</u> (*msh2*Δ), and <u>SRP152953</u> (*zip3*Δ). Scripts, tools, Software and additional data are available at: https://github.com/Neale-Lab and https://github.com/NealeTools/RecombineSim

## Methods

### Yeast Strains

All Saccharomyces cerevisiae strains used in this study are derivatives of SK1 (Kane and Roth, 1974) and S288c (Mortimer and Johnston, 1986). Hybrid strains, utilised in genome-wide mapping, were derived from a cross of haploid SK1 and S288c, or used published datasets from a cross of S96 × YJM789 (Chen et al., 2008; Mancera et al., 2008; Oke et al., 2014; Al-Sweel et al., 2017). Strain genotypes are detailed in (**Supplementary Table 2**). Knockouts were performed and tested by standard transformation and PCR techniques (Longtine et al., 1998). *msh2*Δ::*kanMX6* and *zip3*Δ::*HphMX* were generated by PCR mediated gene replacement using a pFA6a-*kanMX6* or pFA6-*hphMX* plasmid (Goldstein and McCusker, 1999). The *P_GAL_-NDT80::TRP1* allele has the natural *NDT80* promoter replaced by the *GAL1-10* promoter, and strains include a *GAL4::ER* chimeric transactivator for β-estradiol-induced expression (Benjamin et al., 2003). For cytological analyses in hybrid strains, Zip1-GFP (White et al., 2004) was expressed heterozygously from the SK1 parent only. S288c × SK1 hybrids create viable spores (91.98% WT, 72.99% *msh2*Δ spore viability), limiting observational bias that may arise from assaying a limited, surviving population (Crawford et al., 2018).

### Meiotic Timecourse (*ndt80AR* strains)

Diploid strains were incubated at 30°C on YPD plates for 48h. For SK1 diploids, a single colony was inoculated into 4 mL YPD (1% yeast extract, 2% peptone, 2% glucose) and incubated at 30°C at 250 rpm for 24 h. For hybrid crosses, haploid parental isolates were mated in 1 mL YPD for 8 h. An additional 3 mL of YPD was subsequently added and the cells were grown for 16 h. Cells were inoculated to a density of (OD600) 0.2 into 30 mL YPA (1% yeast extract, 2% peptone, 1% K-acetate) and incubated at 250 rpm at 30°C for 14h. Cells were collected by centrifugation, washed in H_2_O, and resuspended in 30mL pre-warmed sporulation media (2% potassium acetate, 5 μg/mL Adenine, 5 μg/mL Arginine, 5 μg/mL Histidine, 15 μg/mL Leucine, 5 μg/mL Tryptophan, 5 μg/mL Uracil). The culture was then incubated at 30°C at 250 rpm for the duration of the time course. After 8h, 2 mL of the synchronised cultures were split and exposed to β-estradiol to a final concentration of 2 mM, which induces the transcription of *NDT80* and thus sporulation. Cultures were then incubated to a total of 48 h at 30°C prior to dissection. For microscopy cells were pre-grown in BYTA medium (buffered 1% yeast extract, 2% tryptone, 1% K-acetate) and *ndt80AR* cultures were not released with β-estradiol.

### Tetrad Dissection

In order to produce hybrid spores for sequencing, SK1 × S288c haploid parents were mated for 8-14 h on YPD plates, with the exception of *ndt80AR* strains, which were mated and grown in liquid YPD for 24 h (see above). Haploids were mated freshly on each occasion and not propagated as diploids in order to reduce mitotic recombination. Sporulation was induced, and tetrads were dissected after 72 h in 2% potassium acetate. For octads, spores were additionally grown for 4–8 h on YPD plates until a single mitotic division had completed, after which the mother-daughter pair were separated. Colonies were grown for 16 h within liquid YPD for genomic DNA extraction. Only tetrads and octads producing four or eight viable spores/colonies, respectively, were considered for genotyping by NGS.

### NGS Library Preparation

Genomic DNA was purified from overnight, saturated YPD cultures using standard phenol-chloroform extraction techniques. Samples of genomic DNA were diluted to 0.2–0.3 ng/μL. DNA concentration was measured using the Qubit High Sensitivity dsDNA Assay. Genomic DNA was fragmented, indexed and amplified via the Nextera XT DNA library Prep Kit according to the best practices recommended by Illumina. In order to check fragment-length distribution and concentration of purified libraries, 1 μL of undiluted library was run on an Agilent Technology 2100 Bioanalyzer using a High Sensitivity DNA chip. To pool samples for sequencing, 5 μL of each sample was combined into a 1.5 mL tube and mixed. 24 μL of the mix was transferred to a tube containing 570 μL hybridisation buffer. The mix was boiled at 96°C for 2 minutes and placed in ice water for 5 minutes. 6 μL of denatured PhiX control (prepared according to Illumina protocol, final concentration 1%) was added to the library, mixed well and then loaded into a MiSeq reagent cartridge. Sequencing was performed in-house using an Illumina MiSeq instrument.

### Alignment, SNP and indel Detection

Individual spores were sequenced to an average read-depth of ~45x. Initially, paired-end read FASTQ files are aligned, via Bowtie2 (Langmead and Salzberg, 2012), to the SacCer3 reference genome (v. R64-2-1; (Engel and Cherry, 2013)) using the parameters: -X 1000 —local —mp 5,1 -D 20 -R 3 -N 1 -L 20 -i S,1,0.50. In order to create a custom SK1 genome to facilitate more accurate genotype-calling, SNP and indel polymorphisms were detected using the GATK (GenomeAnalysisToolkit) function *HaplotypeCaller* (Van der Auwera et al., 2013). An in-house script (*VariantCaller.pl*) subsequently parses the resulting VCF files from 72 spores to calculate: (i) the call frequency (% of spores any given allele is present within), (ii) the cumulative allelic read depth (% of reads that contain a specific allele at a specific loci), and (iii) the cumulative total read depth. To identify legitimate SNPs and indels, variants were filtered for a call-frequency between 44-55%, a total read depth of >250 and an allelic read depth of 95%. Variants within repeat regions, long terminal repeats, retrotransposons and telomeres were also discarded—yielding a final, robust list of 64,591 SNPs and 3972 indels amounting to ~0.57% divergence. A custom SK1 genome (SK1_Mod) was then generated by modifying SacCer3 (v. R64-2-1) to include all filtered/called SNPs and indels.

### Genotype-Calling

Spore data from individual samples was aligned to both the custom SK1_Mod genome and the SacCer3 reference (see below). Alignment produced a SAM file, which was converted into a sorted BAM file using the Samtools function, *view* (Li et al., 2009), for downstream processing. The PySamStats (v. 1.0.1, Miles & Mattioni) module, *variation*, was used to process the sorted BAM file for each sequenced spore, producing a list of the number of reads containing A/C/T/G, insertion or deletion for each genomic position specified in the S288c and SK1 references. Variant reads were isolated and genotyped using in-house, custom scripts as follows. Genotypes were assigned according to the rules: (i) A minimum coverage-depth of 5; (ii) A SNP was called as having the variant genotype if >=70% of the reads at that position match the called variant, or as reference if =>90% of the reads match the reference; (iii) If the variant and reference reads were above 90% of all reads and within 70% of each other, the position was called as heteroduplex; (iv) indels are called as having the variant genotype if >=30% of the reads at that position matched the variant. Such a low threshold was utilised because alignment of indel sequences is biased towards the reference, which means that they are unlikely to be erroneously called as matching the variant genotype. For an indel to be called as the reference genotype, >=95% of the reads must match the reference sequence and there must be fewer than two reads matching the variant call. Any variants that fall below these thresholds were discarded. Genotype calls were converted into a binary signal, either 1 for S288c or 0 for SK1.

### Event Calling

Using the binarised input, chromosomes were split into segments with the same segregation pattern using published scripts (Marsolier-Kergoat et al., 2018). Segment types (i.e. 1:7, 2:6, 2:6, 3:5, 4:4, 4:4*, 5:3, 6:2, 6:2* or 7:1 as previously described (Martini et al., 2011; Marsolier-Kergoat et al., 2018) were also recorded. Recombination events were subsequently called as being a set of segments located between two 4:4 segments longer than 1.5 kb (Marsolier-Kergoat et al., 2018). A 4:4 segment corresponds to a Mendelian segregation profile, 5:3 and 3:5 segments to half-conversion tracts, and 6:2 and 2:6 segments to full conversion tracts. Each recombination event can contain between 0–2 COs or NCOs. Events were additionally classified by the number of chromatids involved (i.e. 1, 2, either sister or non-sister, 3, 4). To ensure compatibility with our data-analysis pipeline, published binarised input data (“segFiles’) from the S96 × YJM789 hybrid (Chen et al., 2008; Mancera et al., 2008; Oke et al., 2014; Al-Sweel et al., 2017), were minimally processed to match column naming, with spores from tetrads each duplicated to create a fake “octad”. This conversion involved no changes in data, only minimal reformatting. Recombination events were then called in the same way as for SK1 × S288c octads generated for this study. Any small differences in CO and NCO counts and positions between the resulting data and that published are likely, therefore, to be due to subtle differences in the event calling criteria used (for example event merging thresholds).

### Event Position & Inter-Crossover Distances (ICDs)

Crossover position, or “midpoint”, is defined as the distance between the mid-points of the first and last SNP/indel markers—an estimate of true event tract length. Inter-crossover distances (ICDs) were then calculated as the distance (in bp) between successive CO midpoints.

### (γ)-Mixture Modelling

Distributional analysis of CO distributions is complicated by the existence of non-interfering, Mus81-Mms4 class II COs—indistinguishable from interfering ZMM-dependent class I COs in our assay. In essence, meiotic ICDs represent a heterogenous, mixed system (**Fig. 2b**) with unknown quantities of each subclass. Latent variables (e.g. class II CO %) may, however, be inferred through probabilistic and statistical methods. Expanding upon the use of the gamma (γ) distribution to model meiotic ICDs from this type of data (Chen et al., 2008; Anderson et al., 2015), experimental data was deconvoluted by fitting two (γ) distributions—one for each subclass of CO—via an expectation maximisation (EM) algorithm (MATLAB 2018a). EM is a commonly applied method for iterative clustering and parameter estimation in mixed models (Do and Batzoglou, 2008).

Briefly, any given ICD was assigned a probability reflective of how likely it is to belong to one of the two sub-distributions. Subsequently, sub-distributions were iteratively shifted, and data point identity was reassigned until a maximum likelihood (ML) solution was converged upon. One (γ) distribution is expected to yield a final (γ)(α) value of ~1.0 (class II, random), while the other is expected to produce a (γ)(α) value of >1.5 (class I, non-random), with their relative contributions to the overall mixture (i.e. the class I:class II ratio) is dependent upon genotype. To validate this approach, simulated ICD datasets of two component mixtures with known parameters, at variable sample sizes (S), were generated using *RecombineSim* and deconvoluted (**Supplementary Fig. 4b**; see below). As a measure of accuracy, the average % difference (N(%Δ)) between estimated and actual parameters was calculated. To calculate average percentage differences N(%Δ) generated by the simulation, each simulated mixture (at each sample size) was performed 100 times. Gamma-parameter estimates were obtained for each simulation. The parameters obtained each time were: γ(α)1, γ(β)1, γ(α)2, γ(β)2, and the proportional weight of the non-random gamma present in the mixture (W1). Estimated values of each parameter were then compared to the actual values used to generate the simulated mixture via the standard percentage difference formula: (|X1-X2| / [(X1+X2)/2])*100. This calculation was repeated for all five of the estimated parameters, with each averaged across the 100 repeats to obtain the final estimated percentage difference value for each parameter. Because final errors for each of the five parameters were found to be similar to one other within each trial, they were averaged to obtain a single N(%Δ) value (**Supplementary Fig. S4b-c**). Accuracy is dependent upon sample size (S) and to a lesser extent on the relative proportions of each subpopulation—and thus how likely a subpopulation is to be readily observed within the mixed population. For example, (γ) mixtures containing 10 or 25% class II COs exhibit average errors of 10.0% and 9.1% at (S) = 500 and 4.53% and 3.73% at (S) = 2000 respectively (**Supplementary Fig. 4c**). Experimental datasets range from (S) values of 354 to 3365, therefore reasonable error rates of ~<10% were expected.

A similar mixed-modelling approach (CODA) has been used to estimate relative proportions of mixed gamma populations for distributions of COs along multiple observations of a chromosome (Gauthier et al., 2011). Initial attempts to use this software failed presumably due to our low sample number. We circumvented this problem by concatenating the genome-wide set of inter-CO distances within a single meiosis to generate single pseudo (giant) chromosome encompassing, effectively, the entire yeast genome. Best-fit non-random gammas and proportions of the non-random (sprinkling) component were subsequently estimated as: Wild type (alpha = 3.4, proportion 0.28); *msh2*Δ (alpha = 3.9, proportion 0.11). Such estimates are similar to those obtained using our direct gamma mixture-modelling algorithm (e.g. **Fig. 2d**).

### Simulating patterns of class I and class II CO formation

Randomised or mixed (class I + class II) ICD simulations were performed using a simulation platform (*RecombineSim*) built in MATLAB 2018a. A typical simulation run is depicted in (**Supplementary Fig. 4a**). In brief, virtual *S. cerevisiae* chromosomes are constructed as binned, numerical arrays at a 100 bp resolution adjusted to reflect the limit of experimental detection governed by the leftmost and rightmost genetic markers (SNPs/indels). Any given 100 bp bin possesses a numerical recombination potential (recom(P)), which governs the ability of an interfering CO to successfully form at that site. Once formed within the simulation, class I COs impose a zone of “interference”, by altering recom(P) values in adjacent bins in a distance-dependent manner—a similar principle to the beam-film model of CO interference (Zhang et al., 2014a). The exact shape and width of interference imposed was determined by the best fit (γ) parameters (α, β) obtained from the MLE mixture modelling (see above) for the genotype currently being simulated—and applied as a hazard function (EQN 1.1):

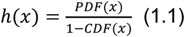

A hazard function describes the probability that, given a pre-existing CO at position x(0), another CO will form at any given distance (x) away (Chen et al., 2008)—and thus is a natural representation of interference. A fractional amount of class II COs that remain insensitive to recom(P) are introduced via the C_PROB_ parameter where necessary as in gamma-sprinkling models (Copenhaver et al., 2002; Housworth and Stahl, 2003; Housworth and Stahl, 2003; Falque et al., 2009). In order to closely match the in vivo datasets, simulated CO events arising within 1.5 kb of one another were also merged, creating a single visible event at the midpoint.

To explore the impact that failed class I CO maturation may have on genome-wide patterns, simulations were additionally performed in which, for a controlled (and variable) fraction of class I events, the class I event itself was removed from the population of counted events only after implementation of interference around the site of this precursor event. Such “failed-maturation” events are thus influenced by pre-existing patterns of interference, and indeed influence the probability of flanking class I CO events that may arise later in the simulation, but are not themselves counted, and thus do not contribute directly to the final pattern or frequency of CO events reported.

In all cases, simulations (N = 10,000 cells) were iterated until the final frequency of visible COs (class I plus class II) per simulated cell equaled the frequency observed for a given genotype. Thus, when events were merged, and/or when events were removed (to simulate maturation failure), additional CO events were simulated for those cells.

### ICD Transformation

The formation of a variable number of events (N) within a finite space (*lim*) (i.e. a chromosome or genome length) skews CDFs i.e. a higher CO frequency causes a downward shift in ICD size. An ICD distribution produced under identical spatial rules but with a different event count would therefore generate significantly different CDFs—failing or biasing statistical testing and undermining the ability to assess distributional agreement. This skew can be readily observed using simulated data (**Supplementary Fig. 1a**). Notably, higher values of (N) cause a leftward skew. The relationship between (N) and ICD size for a given *lim* is, however, linear (Batten et al., 1993). Consequently, in order to isolate the distributional identity of any given sample (i.e. isolate γ(α) from γ(β)), ICD data can be transformed by calculating the product of ICD size (ICD × event count). Data transformation results in perfectly aligned CDFs despite varying (N), validating this approach (**Supplementary Fig. 1b**).

### Statistical Analyses

A *Kolmogorov-Smirnov* goodness-of-fit (GoF) test is a non-parametric test used to compare continuous probability distributions in order to assess the null hypothesis that both samples derive from identical populations, based on their maximal difference (D_KS_) (Massey Jr, 1951; Miller, 1956). (P) values of the KS-test effectively describe the probability that, if the null hypothesis is true, the observed CDFs would be as far apart as observed. (P) values may therefore constitute an indirect measure of distributional agreement, as employed throughout this paper. KS-tests were performed using the MATLAB 2018a packages: *kstest* and *kstest2*. A two-sample T-test was utilised to determine whether a difference in mean value is significant or has arisen by chance. Two-sample T-Tests were performed using the MATLAB 2018a package: *ttest2*.

### Microscopy and Cytological Analysis

4.5 mL of meiotic culture was spun down on a bench centrifuge and resuspended to 500 μL with 1M pH 7.0 D-Sorbitol. 12 μL of 1.0 M DTT and 7 μL of 10 mg/mL Zymolyase in 10% glucose solution was added and cells were spheroplasted by incubation at 37°C for 35–50 min with agitation. Spheroplasting success was determined by taking 2–3 μL of the solution and adding an equivalent volume of 1.0% (w/v) Sodium N-Lauroylsarcosine while under microscopic observation. Cells should immediately lyse as the exposed membrane is disrupted by the detergent. 3.5 mL of Stop Solution (0.1M MES, 1 mM EDTA, 0.5 mM MgCl_2_, 1M D-Sorbitol, pH 6.4) was subsequently added and the cells were spun down to be resuspended in 100 μL Spread Solution (0.1M MES, 1 mM EDTA, 0.5 mM MgCl_2_, pH 6.4) and distributed between four slides, which had been soaked in 70% ethanol overnight and wiped clean. To each slide, fixative (4.0% (w/v) formaldehyde, 3.8% (w/v) Sucrose, pH7.5) was added dropwise, followed by detergent (1% Lipsol, 0.1% Bibby Sterilin) to a ratio of 1:3:6 (suspension : fixative : detergent) before lightly mixing and incubating for 1 min at room temperature (RT). Further fixative was added dropwise to a final ratio of 1:9:6 and the mixture spread across the slide. Each spread was subsequently incubated at RT for 30 min in damp conditions, then allowed to air dry at RT overnight. Once dry, slides were sequentially washed in 0.2% (v/v) PhotoFlo Wetting Agent (Kodak) and dH_2_O, and stored at 4°C.

Slides were washed once in 0.025% Triton X-100 for 10 min at RT and twice in PBS for 5 min at room temperature. Slides were blocked in 5% skimmed milk with PBS for 3 h at 37°C. Excess liquid was removed and slides laid horizontally in damp conditions. 40 μL of primary antibody (anti-Zip3 (Shinohara et al., 2008) from rat at 1:200 and/or anti-Red1 (Genecust, affinity purified, raised against aa(426-827)) from rabbit at 1:200) in 1% skimmed milk with PBS was added under coverslips. Slides were incubated at 4°C overnight (15.5 h) and washed three times in PBS for 5 min at RT. Excess liquid was then removed and slides were returned to damp conditions. 40 μL of secondary antibody (anti-rat AlexaFluor555 at 1:200 and anti-rabbit AlexaFluor488 at 1:500) in 1% skimmed milk with PBS iwas added under coverslips. Slides were incubated at room temperature for 2.5 h and then washed three times with PBS for 5 min at room temperature. Cover slips were affixed using Vectashield mounting medium with DAPI, sealed with clear varnish and imaged on an Olympus IX71 (z = 0.2 μM, Exposure times: TRITC-mCherry = 0.2 sec, eGFP = 1.0s, DAPI = 0.1s). Images were randomised, deconvoluted via Huygens (software) and foci were automatically counted using an in-house plugin for ImageJ (FindFoci) as previously described (Herbert et al., 2014), with an appropriate mask to discard signals outside of nuclei. For Zip3 interfoci-distance scoring, pixels denoting the centre of each Zip3 focus, and Zip1 ends, were manually selected along clearly separable bivalents as determined by Zip1-GFP signal (120-156 bivalents per strain, error margin of approximately 1 pixel = 0.1 µm). These positions were selected using the ImageJ segmented line tool and segment lengths were then calculated by macro, confirming agreement with total length as measured by ImageJ standard tool.

## Supplementary Discussion

### The *SK1-ML3* allele has a reduced capacity to generate CO interference

As previously noted, wild type CO frequencies are higher within S96 × YJM789 than S288c × SK1 (91.4 vs 74.3 COs/meiosis) (**Fig. 1b**; **Supplementary Table 1**). Moreover, CO distributions deviate even further from that expected for a random distribution in S96 × YJM789 (p<0.01; Two-sample KS test) (**Supplementary Fig. 2a**) consistent with the (γ)-mixture modelling results (**Fig. 2d**) suggesting that the class I CO fraction is greater in S96 × YJM789 (75% vs. 67%). Molecular incompatibilities between certain alleles of the CO formation or CO interference machinery may account for these cross-specific differences (Al-Sweel et al., 2017). To investigate this hypothesis further, we analysed the frequency and distribution of COs within an *mlh3*Δ S288c × YJM789 background containing ectopically expressed copies of the wild-type SK1 *MLH1* and *MLH3* alleles (*SK1-MLH3*) (Al-Sweel et al., 2017). Remarkably, introduction of the SK1 alleles was sufficient to alter the genome-wide pattern of COs, producing a relative distribution identical to that observed in the S288c × SK1 hybrid (p = 0.91; Two-sample KS Test) (**Supplementary Fig. 2b**). Surprisingly, despite this change in CO distribution, introduction of the SK1 alleles did not significantly reduce CO frequency when compared to the large pool of 51 wild-type S96 × YJM789 tetrads (90.9 COs vs 91.4 per meiosis) (**Supplementary Table 1**). However, the observed frequency was significantly reduced relative to the subset (n=5) of S96 × YJM789 tetrads generated independently (100.4 vs. 90.9 COs per meiosis; p < 0.01; Two-sample T-test; (Chen et al., 2008)). Collectively, these results suggest that, in some manner, the SK1 *MLH1* and/or *MLH3* alleles are partially deficient in class I CO formation, and/or propagation and/or sensitivity to CO interference. However, the much lower CO frequency observed within S288c × SK1 hybrids (74.3 COs per meiosis) cannot be fully ascribed to the SK1 *MLH1* and/or *MLH3* alleles alone. In the context of this study, the relative inefficiency of class I CO formation in the SK1 × S288c hybrid enhances the visibility of the *msh2*Δ-induced CO phenotype relative to in the S96 × YJM789 hybrid (**Fig. 1c-d**).

### Zip3 foci located at the termini of Zip1-stained synapsed axes

We noted that within our microscopic analysis of synapsed chromosomes a large proportion of Zip3 foci appeared to occur at the terminal ends of Zip1 stretches (**Fig. 3c-d**). However, we suggest that this is less likely to represent disproportionate Zip3 occupancy at chromosome ends than the presence of at least some fraction of the chromosome length (i.e. telomere proximal) that is not visible in this assay—perhaps suggesting that Zip1 loading and/or polymerisation beyond the most terminal Zip3 focus is inefficient and/or destabilised during the spreading procedure. Indeed, in a similar analysis, Zhang *et al* (Zhang et al., 2014b) observed telomeric LacO/LacI-GFP staining (on Chr. XV) beyond the end of the observable synaptonemal complex as stained by Zip1. Such observations are also consistent with the observations that chromosome ends disproportionately retain markers of incomplete synapsis and persistent DSB formation (Subramanian et al., 2019), and have differential compaction that is Zip1-dependent (Schalbetter et al., 2019) at this *ndt80*Δ-induced pachytene-arrest stage.

### Localised impact of polymorphism density upon CO formation

S288c × SK1 variants have an average density of 1 per 175 bp and a median inter-variant distance of 81 bp (93.12% of inter-variant distances are <500bp) and are therefore evenly spaced and present at high density across each chromosome. Maps of polymorphism density are shown in (**Supplementary Fig. 7c-d**) for two example chromosomes. In general, S288c × SK1 chromosomes are organised into local peaks and troughs of variant density while maintaining overall uniformity. Therefore, it seems unlikely that inactivation of Msh2 would result in a gross-redistribution of CO formation toward any particular region of the chromosome as it may do within organisms with less uniform SNP/indel density, such as *A. thaliana* (Ziolkowski et al., 2015). To further investigate the way in which polymorphisms sculpt the meiotic landscape, we repeated the analysis shown in (**Fig. 4b-c**) using expanded ±1000 bp and ±2000 bp windows (**Supplementary Fig. 7a-b**). A Msh2-dependent and statistically significant skew toward regions of lower sequence divergence (p<0.01; Two-sample KS test) is retained at ±1000 bp but is significantly diminished at ±2000 bp (p=0.51; Two-sample KS test), suggesting that mismatched sequences have the greatest impact when present within the recombination intermediate structures.

## Supplementary Figures

**Supplementary Fig. 1.**
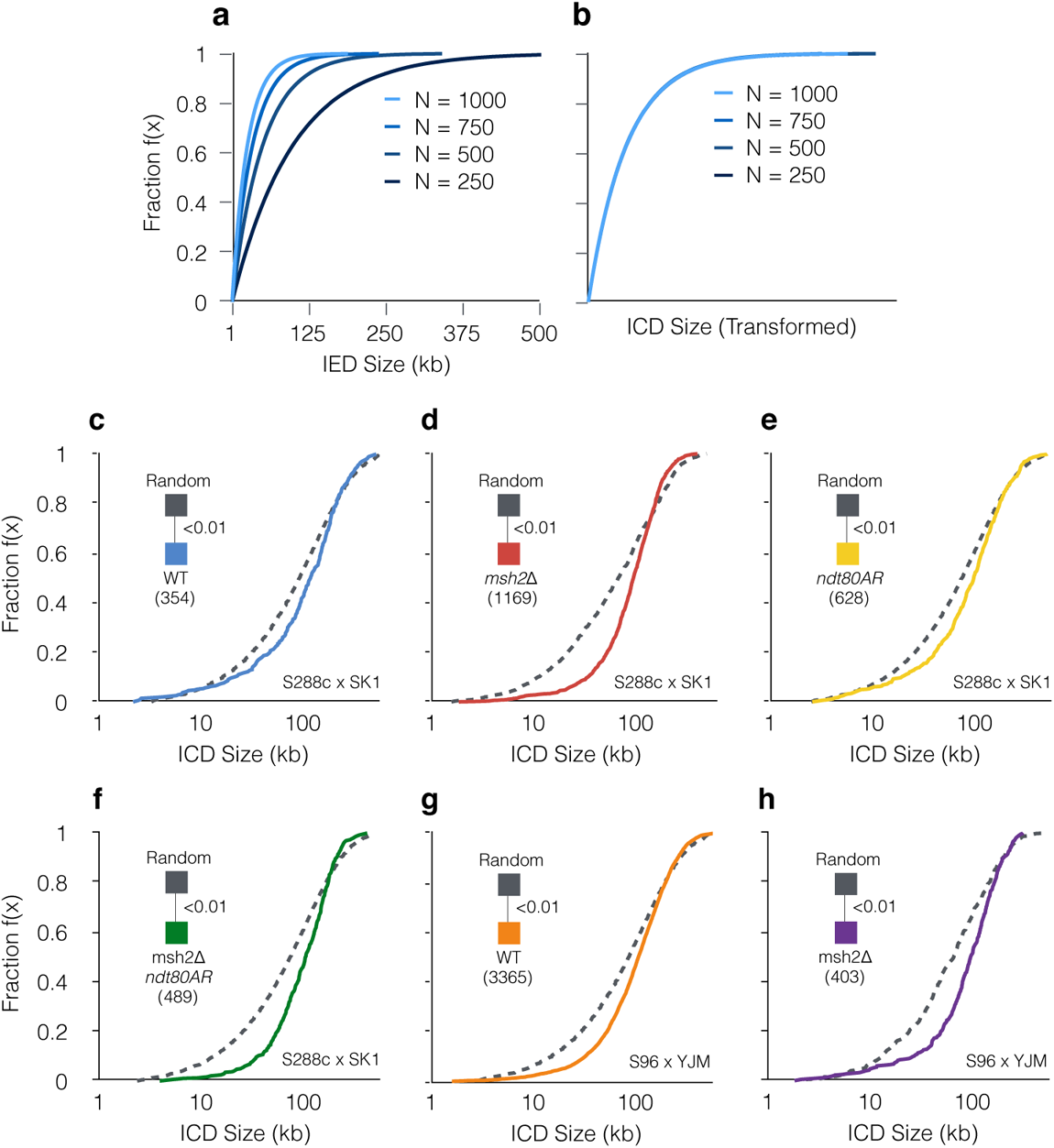
CO interference is present to varying degrees within all mapped strains. **a,** Empirical cumulative distribution function (eCDF) showing ICD data derived from interfering simulations (γ(α) = 3.0) at varying CO per cell frequencies (N). **b,** As in **(a)** but ICDs are transformed (see **Methods**) to correct for skews generated by differing CO frequencies. **c-h**, eCDFs showing the fraction of ICDs at or below a given size. The total number of experimental ICDs is indicated in brackets. Randomised datasets were generated via simulation to represent a state of no interference (**Methods**). Pairwise goodness-of-fit tests were performed between genotypes as indicated (triangular legend). P values: Two-sample KS-test.

**Supplementary Fig. 2.**
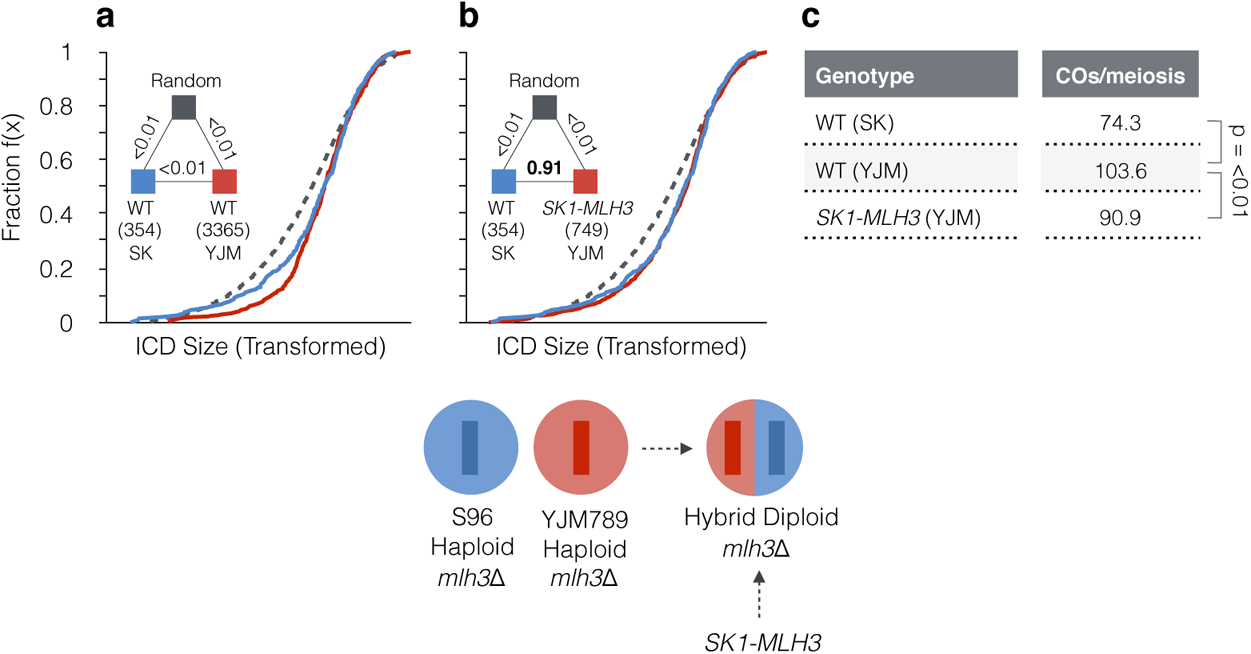
Cross-specific differences—the *SK1-MLH3* allele has reduced capacity to mediate CO interference. **a-b,** Empirical cumulative distribution functions (eCDFs) showing the fraction of ICDs at or below a given size. The total number of experimental ICDs is indicated in brackets. ICDs are transformed (see **Methods**) to correct for skews generated by differing CO frequencies. Randomised datasets were generated via simulation to represent a state of no interference (**Methods**). Pairwise goodness-of-fit tests were performed between genotypes as indicated (triangular legend). A schematic of the *SK1-MLH3* strain analysed is shown. P values: Two-sample KS-test. **c,** Average number of COs per meiosis for each genotype. P values: Two-sample T-test.

**Supplementary Fig. 3.**
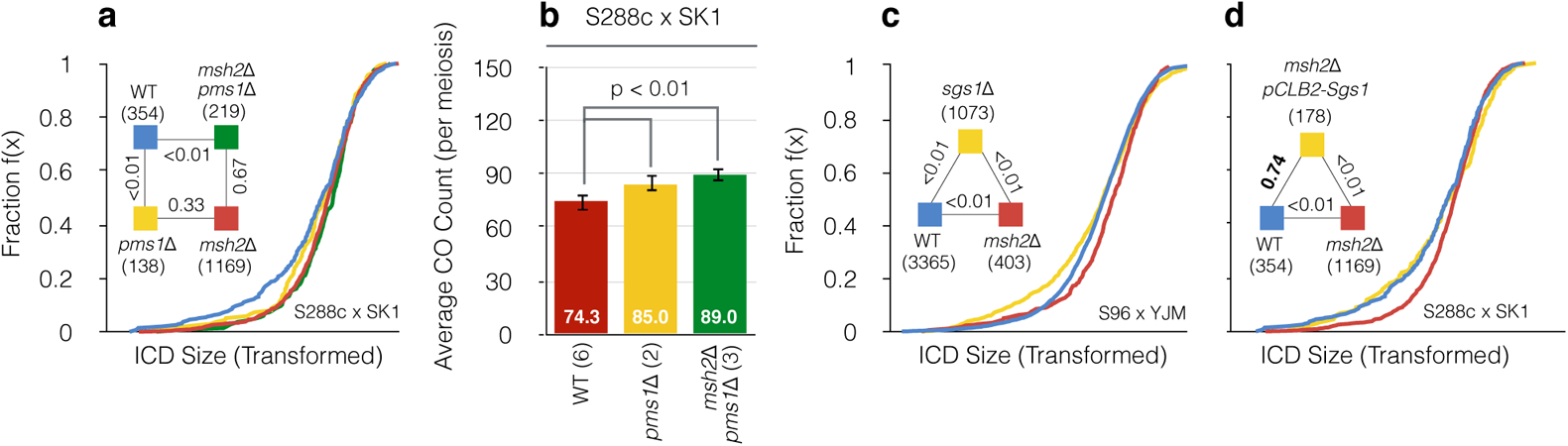
Mechanistic details of MMR-dependent suppression of interfering COs. **a,** Empirical cumulative distribution function (eCDF) showing the fraction of ICDs at or below a given size. The total number of experimental ICDs is indicated in brackets. ICDs are transformed (see **Methods**) to correct for skews generated by differing CO frequencies. Randomised datasets were generated via simulation to represent a state of no interference (**Methods**). Pairwise goodness-of-fit tests were performed between genotypes as indicated (triangular legend). P values: Two-sample KS-test. **b,** Average number of COs per meiosis for each genotype. The number of individual meioses sequenced per genotype is indicated. Error bars: 95% confidence intervals (CI*). P values: Two-sample T-test.* **c-d,** As in **(a)** but for differing genotypes.

**Supplementary Fig. 4.**
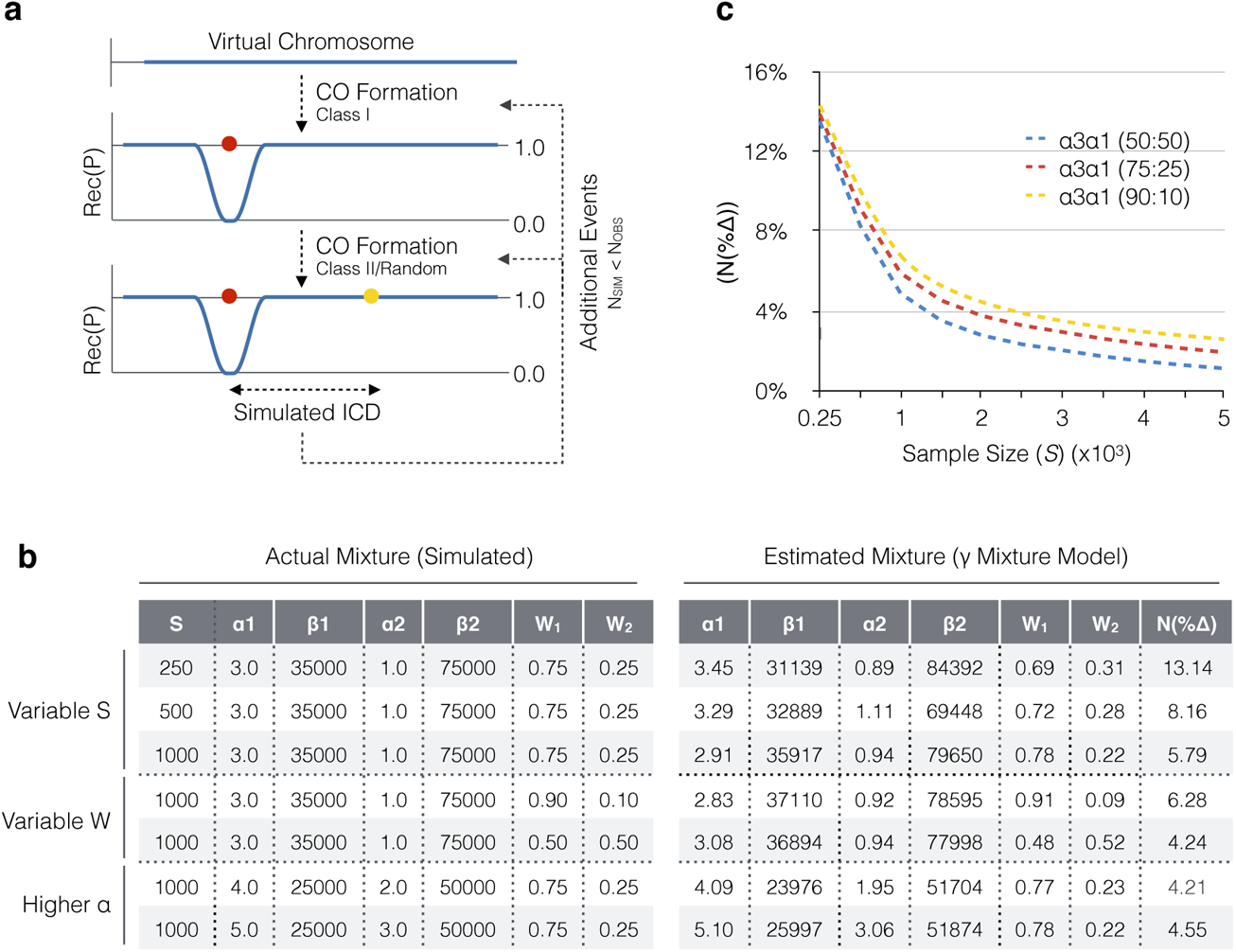
Modelling CO distributions. **a,** *RecombineSim* overview. Virtual chromosomes are constructed at a 100bp resolution as binned, numerical arrays upon which meiotic CO formation is simulated (Methods). Any given 100bp contains a value in the range of [0.0-1.0], designating its recombination potential (Rec(P)). Prior to CO formation, bins are initially populated with [1.0]—denoting an equal probability of class I CO formation in any given bin. During the formation of an interfering CO, *RecombineSim* imposes CO interference as a distance-dependent zone of repression by modifying Rec(P) values according to a hazard function derived from a manually specified γ(α _I_) value, or a γ(α _I_) value estimated from experimental data following gamma (γ) mixture modelling using maximum likelihood expectation (MLE; Methods). Such localised repression around each sequential event thus has the potential to influence the position of all subsequent interfering COs that are simulated. Non-interfering, class II COs are distributed randomly independently of Rec(P) and do not impose, nor are sensitive to, simulated CO interference. Successive events falling within a set threshold of one another (e.g. 1.5 kb) are merged into a single event residing at the midpoint position. These processes repeat until a pre-determined number of simulated ICDs are obtained. **b,** To estimate accuracy of the MLE mixture modelling algorithm, it was used to resolve and estimate individual components of simulated two component mixtures with known parameters (α,β), at known weights (W)—generated via *RecombineSim*. A set of representative examples are shown. Percentage differences between actual and estimated parameters are calculated and averaged to estimate error rate (N(%Δ)) and algorithm accuracy. S = number of ICDs. **c,** Error rate (N(%Δ)) values for three (γ) mixtures calculated at varying sample size (S).

**Supplementary Fig. 5.**
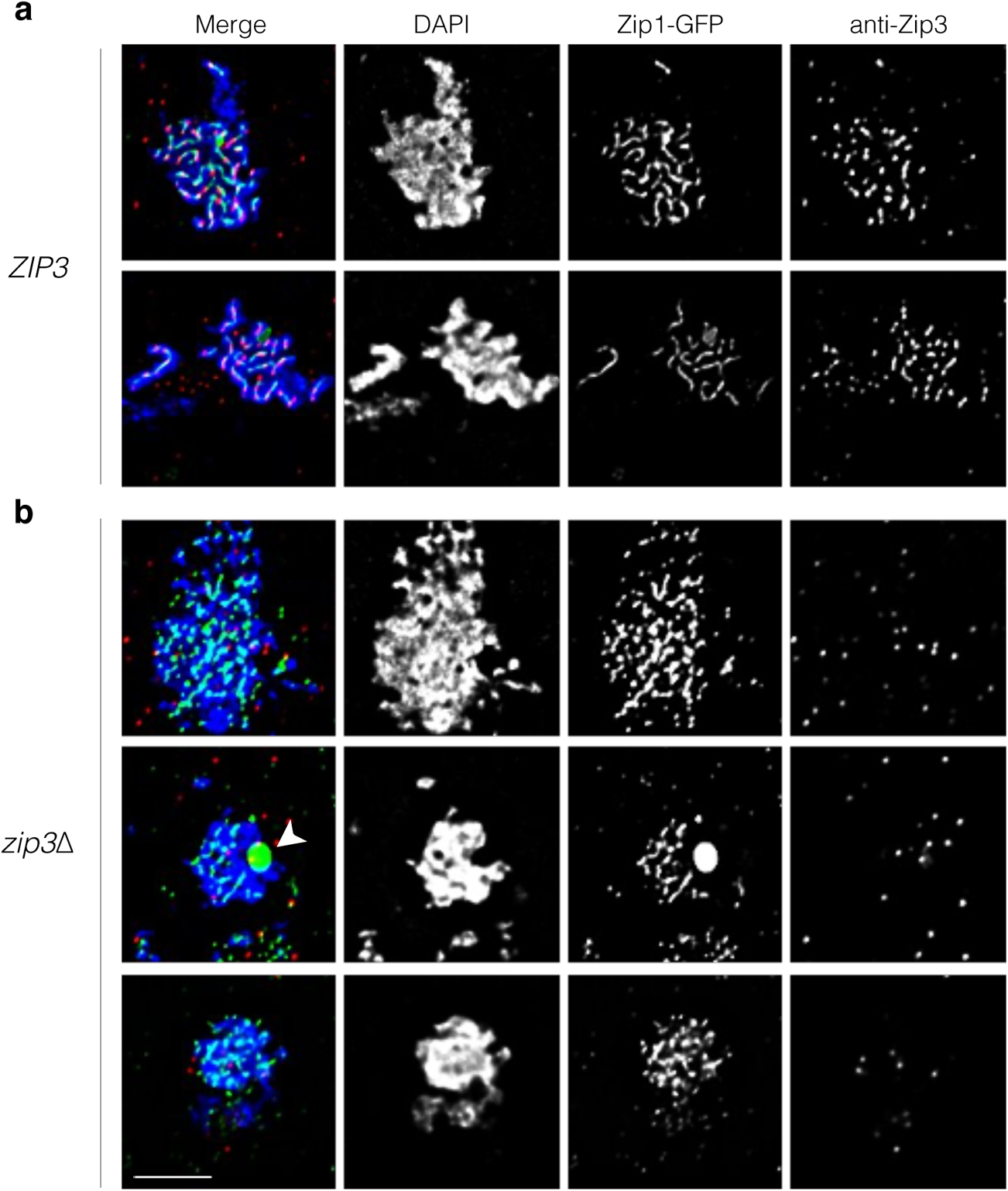
Specificity of anti-Zip3 antibody. **a-b,** Representative chromosome spreads of control (**a**) and *zip3*Δ (**b**) cells at the approximate pachytene stage of meiosis, indicated by Zip1-GFP thread-like signals in control cells, but more punctate Zip1-GFP patterns in *zip3*Δ cells (the most complete that they become). Occasional Zip1-GFP polycomplexes were also observed (arrowhead). In *zip3*Δ cells, anti-Zip3 staining detected only background random signals arising from random binding on the slides at locations that were not enriched in the areas of spread chromatin (blue DAPI-stained signals). Scale bar = µm

**Supplementary Fig 6.**
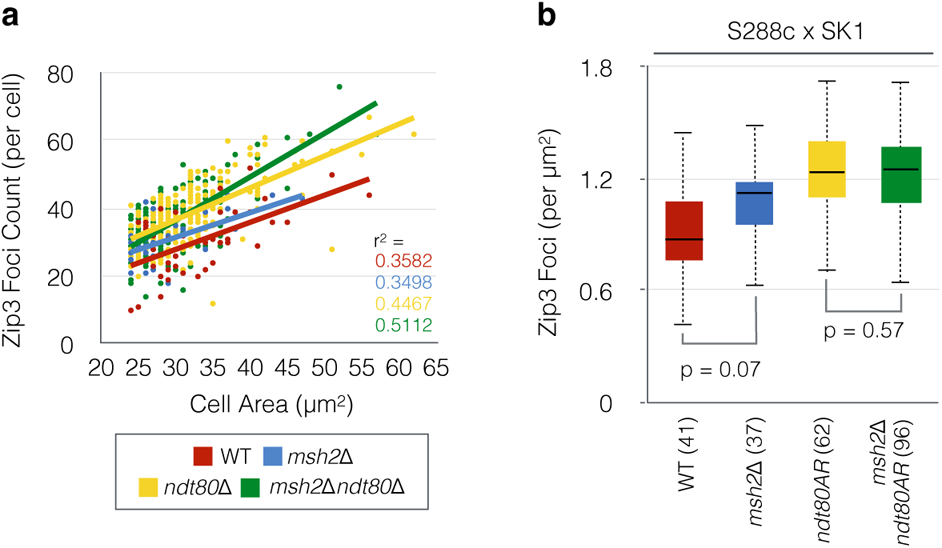
Detected Zip3 foci counts are positively correlated with DAPI-delimited nuclear-spread area. **a,** Scatter plot of Zip3 foci counts per cell against spread area delimited by the DAPI-positive signal for the indicated strains. R-squared correlation values are shown. **b,** Box-and-whisker plot showing Zip3 foci counts per square micron obtained from chromosome spreads of S288c × SK1 *ndt80*Δ cells prepared at 8 h following induction of meiosis (pachytene arrest). Midlines denote median values, box limits are first and third quartile, whiskers are highest/lowest values within 1.5-fold of interquartile range. P values: Two-sample T-test. The total number of nuclei counted is indicated in brackets.

**Supplementary Fig. 7.**
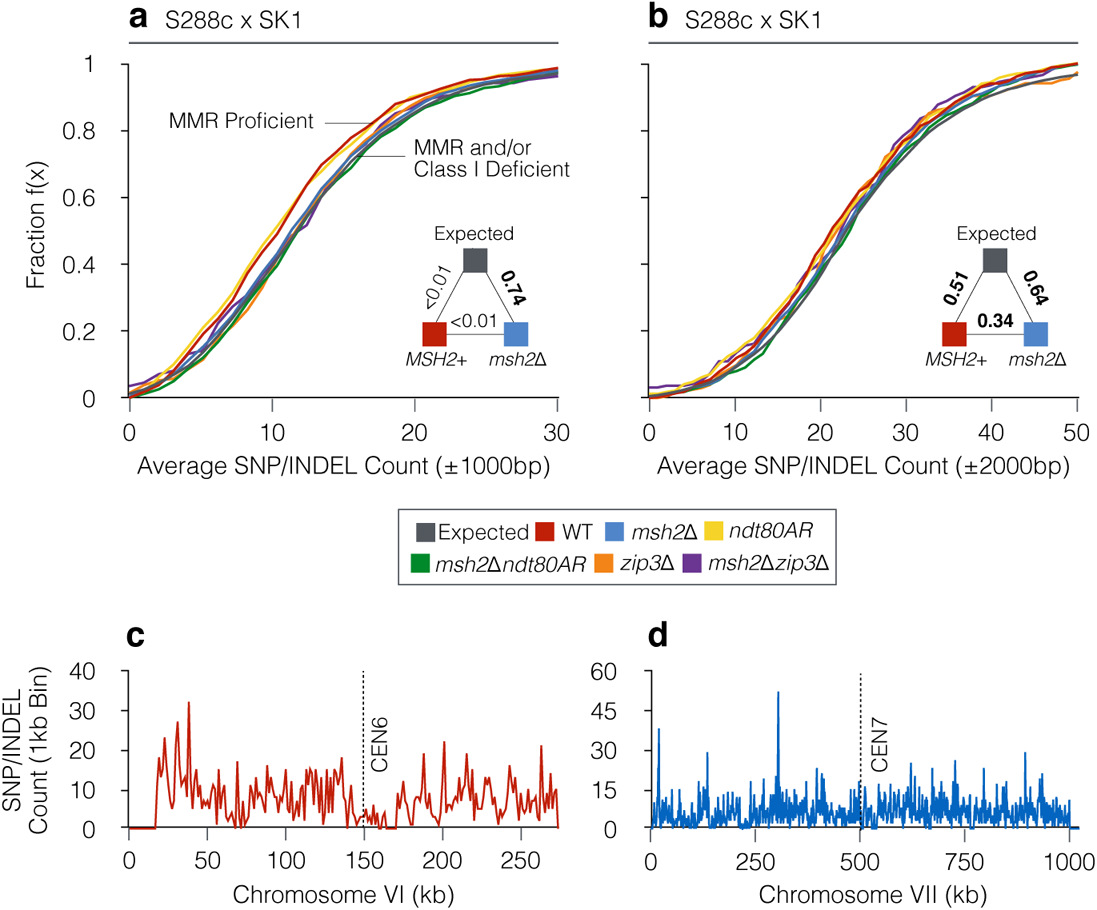
Localised impact of polymorphism density upon CO formation. **a-b,** Empirical cumulative distribution functions (eCDFs) showing the fraction of COs that reside within a region (±1000bp and ±2000bp respectively) of a given SNP/indel count (S288c × SK1 only). Expected is calculated using DSB hotspot midpoints (Pan et al 2011). Pairwise goodness-of-fit tests were performed between pooled *msh2*Δ and *MSH2*^+^ datasets as indicated (triangular legend). P values: Two-sample KS-test. **c-d,** Example smoothed SNP/INDEL density maps per 1 kb bin in the S288c × SK1 hybrid for ChrVI and ChrVII.

**Supplementary Fig. 8.**
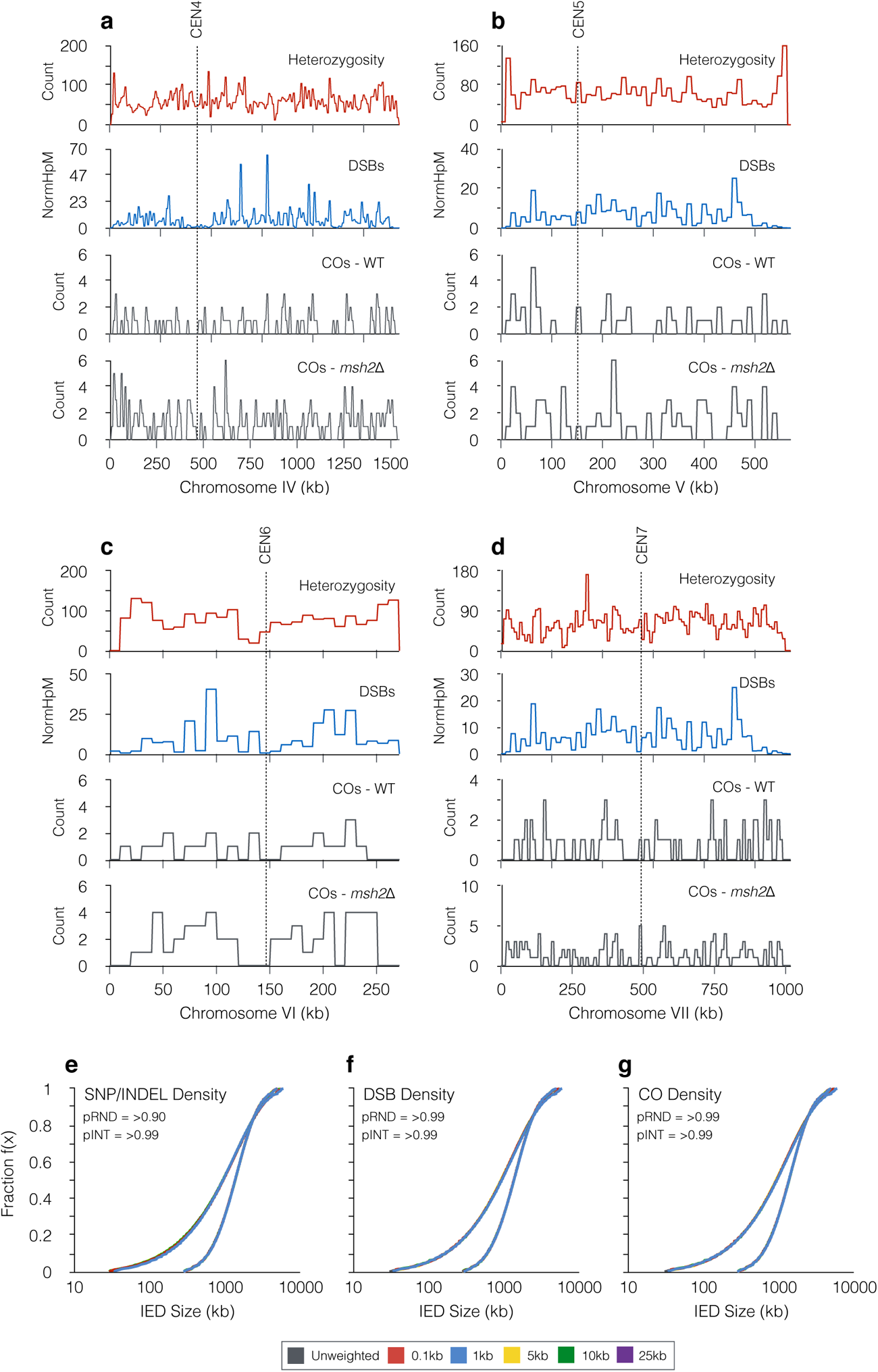
Impact on simulated CO distributions of local deviations in density of heterozygosity, DSBs, and COs. **a-b,** Comparison of spatial distribution of population-average densities of heterozygosity, DSB formation (Pan et al. 2011), and CO formation in wild-type and *msh2*Δ cells for four representative chromosomes binned at 10 kb resolution. Although each chromosome has localised deviation from uniformity, each feature is spread relatively evenly across the length of each chromosome. **e-g,** To test the impact that localised deviations in heterozygosity (**e**), DSB formation (**f**), and observed CO density (**g**) might have on relative distributions of COs, simulations of example random (RND; alpha=1) and interfering (INT; alpha=3) gamma distributions were performed as in **Supplementary Fig. 4**, but additionally weighting CO site selection by the relative amplitude of each parameter at varying levels of smoothing (0.1–25 kb). No change in distributions were observed indicating that the nonuniform distribution of these features is unable to significantly bias relative patterns of CO formation. P values reported are the minimum observed out of the five smoothing values tested for each parameter.

**Supplementary Table 1.** Summary of whole-genome recombination data analysed in this study. Meioses analysed indicate the number of four-spore viable tetrads (or eight-spore viable octads for *msh2*Δ SK1 × S288c derivative strains) analysed. Spore viabilities for the SK1 × S288c strains used in this study (as from Crawford et al 2019) are as follows: Wild type 82.0%, *msh2*Δ 73.0, *ndt80AR* 70.4%, *msh2*Δ *ndt80AR* 73.2%, *zip3*Δ 46.2%, *msh2*Δ *zip3*Δ 35.9%. Other samples analyses employ previously published datasets as indicated by Source column. COs and NCOs are total numbers analysed across all meioses of each genotype, or the average number observed per meiosis. ICDs indicates the total number of inter-crossover distances used to assess CO distributions per genotype. For S96xYJM789 two sources of wild-type data were pooled, as were *P_CLB2_mms4* and *mms4*Δ.

| Relevant genotype | Hybrid | Meioses analysed | Total COs | Total NCOs | COs per meiosis | NCOs per meiosis | ICDs | Source |
| --- | --- | --- | --- | --- | --- | --- | --- | --- |
| Wild type | SK1 x S288c | 6 | 446 | 185 | 74.3 | 30.8 | 354 | Crawford et al 2019; This study |
| *msh2Δ* | SK1 x S288c | 13 | 1377 | 1206 | 105.9 | 92.8 | 1169 | Crawford et al 2019; This study |
| *ndt80AR* | SK1 x S288c | 8 | 756 | 391 | 94.5 | 48.9 | 628 | Crawford et al 2019; This study |
| *ndt80AR msh2Δ* | SK1 x S288c | 6 | 585 | 521 | 97.5 | 86.8 | 489 | Crawford et al 2019; This study |
| *zip3Δ* | SK1 x S288c | 4 | 187 | 344 | 46.8 | 86.0 | 153 | Crawford et al 2019; This study |
| *msh2Δ zip3Δ* | SK1 x S288c | 4 | 104 | 1356 | 26.0 | 339.0 | 75 | Crawford et al 2019; This study |
| *pms1Δ* | SK1 x S288c | 2 | 170 | 234 | 85.0 | 117.0 | 138 | Marsolier-Kergoat et al. 2018 |
| *pms1Δ msh2Δ* | SK1 x S288c | 3 | 267 | 236 | 89.0 | 78.7 | 219 | Marsolier-Kergoat et al. 2018 |
| *P<sub>CLB2</sub>sgs1 msh2Δ* | SK1 x S288c | 2 | 219 | 183 | 109.5 | 91.5 | 178 | Marsolier-Kergoat et al. 2018 |
| Wild type | S96 x YJM789 | 5 | 502 | 261 | 100.4 | 52.2 | 422 | Chen et al 2008 |
| Wild type | S96 x YJM789 | 46 | 4161 | 2128 | 90.5 | 46.3 | 3425 | Steinmetz |
| Pooled wild type | S96 x YJM789 | 51 | 4663 | 2389 | 91.4 | 46.8 | 3365 |  |
| *msh2Δ* | S96 x YJM789 | 4 | 467 | 225 | 116.8 | 56.3 | 403 | Oke et al 2014 |
| *msh4Δ* | S96 x YJM789 | 6 | 214 | 341 | 35.7 | 56.8 | 118 | Oke et al 2014 |
| *zip3Δ* | S96 x YJM789 | 7 | 429 | 852 | 61.3 | 121.7 | 317 | Oke et al 2014 |
| *mms4Δ msh2Δ* | S96 x YJM789 | 4 | 391 | 317 | 97.8 | 79.3 | 327 | Oke et al 2014 |
| *P<sub>CLB2</sub>mms4* | S96 x YJM789 | 3 | 321 | 260 | 107.0 | 86.7 | 273 | Oke et al 2014 |
| *mms4Δ* | S96 x YJM789 | 4 | 406 | 321 | 101.5 | 80.3 | 342 | Oke et al 2014 |
| Pooled *mms4* | S96 x YJM789 | 7 | 727 | 581 | 103.9 | 83.0 | 616 |  |
| *sgs1Δ* | S96 x YJM789 | 11 | 1249 | 848 | 113.5 | 77.1 | 1073 | Oke et al 2014 |
| *SK1-MLH3* | S96 x YJM789 | 10 | 909 | 634 | 90.9 | 63.4 | 749 | Al-Sweel et al 2017 |

**Supplementary Table 2.** Strain Table (S288c × SK1)

| Genotype | Strain | Background | Mat | Genotype |
| --- | --- | --- | --- | --- |
| wild type | MJ513 | SK1 | a | *ho::LYS2 lys2Δ leu2Δ arg4Δ* |
|  | MJ600 | S288c | α | *ade8Δ* |
| *msh2Δ* | MC26 | SK1 | α | *ho::LYS2 lys2Δ ura3Δ arg4 leu2 msh2Δ::Kan* |
|  | MC49 | S288c | a | *ade8Δ msh2Δ::KanMX* |
| *ndt80AR* | MJ43 | SK1 | α | *ho::LYS2 lys2Δ arg4Δ leu2Δ::hisG trp1Δ::hisG his4XΔ::LEU2 nuc1Δ::LEU2 PGAL1-NDT80::TRP1 ura3::pGPD1-GAL4(848)-ER::URA3* |
|  | MC42 | S288c | a | *ade8Δ ndt80Δ::KanMX* |
| *msh2Δndt80AR* | MC298 | SK1 | a | *ho::LYS2 lys2Δ ura3Δ arg4 leu2 trp1Δ::hisG ura3Δ::PGPD1-GAL4(848)-ER::URA3 PGAL1-NDT80::TRP1 msh2Δ::Kan* |
|  | MC300 | S288c | α | *ade8Δ ndt80Δ::KanMX msh2Δ::KanMX* |
| *zip3Δ* | MC322 | SK1 | α | *ho::LYS2 lys2Δ ura3Δ arg4 leu2 zip3Δ::HphMX4* |
|  | MC313 | S288c | a | *ade8Δ zip3Δ::HphMX4* |
| *msh2Δzip3Δ* | MC326 | SK1 | α | *ho::LYS2 lys2Δ ura3Δ arg4 leu2 msh2Δ::Kan zip3Δ::HphMX4* |
|  | MC317 | S288c | a | *ade8Δ msh2Δ::Kan zip3Δ::HphMX4* |
| Wild type | hLH117 | SK1 | a | *ho::hisG, lys2, leu2::hisG, trp1::hisG, his3::hisG, ura3, ZIP1-GFP(at AA700)* |
|  | hLH2 | S288c | α | *ade8* |
| *msh2Δ* | hLH123 | SK1 | α | *ho::hisG lys2, leu2::hisG, trp1::hisG, his3::hisG, ura3, ZIP1-GFP(at AA700), msh2::KanMX* |
|  | MC49 | S288c | a | *ade8 msh2Δ::KanMX* |
| *ndt80AR* | hLH127 | SK1 | α | *ho::hisG, lys2, leu2::hisG, trp1::hisG, his3::hisG, ZIP1-GFP(at AA700), ura3::pGPD1GAL4(848)-ER::URA3, pGAL-NDT80::TRP1* |
|  | MC42 | S288c | a | *ade8Δ ndt80Δ::KanMX* |
| *msh2Δndt80AR* | hLH130 | SK1 | a | *ho::LYS2, lys2, leu2::hisG, trp1::hisG, his3::hisG, ZIP1-GFP(at AA700), msh2::KanMX, ura3::pGPD1GAL4(848)-ER::URA3, pGAL-NDT80::TRP1* |
|  | MC300 | S288c | α | *ade8Δ ndt80Δ::KanMX msh2Δ::KanMX* |

